# Protein language models accelerate the discovery of Plastic-Degrading Enzymes

**DOI:** 10.1101/2025.02.09.637306

**Authors:** David Medina-Ortiz, Diego Alvarez-Saravia, Nicole Soto-García, Diego Sandoval-Vargas, Jacqueline Aldridge, Sebastián Rodríguez, Barbara Andrews, Juan A. Asenjo, Anamaría Daza

## Abstract

Plastic pollution presents a critical environmental challenge, necessitating innovative and sustainable solutions. In this context, biodegradation using microorganisms and enzymes offers an environmentally friendly alternative. This work introduces an AI-driven framework that integrates machine learning (ML) and generative models to accelerate the discovery and design of plastic-degrading enzymes. By leveraging pre-trained protein language models and curated datasets, we developed seven ML-based binary classification models to identify enzymes targeting specific plastic substrates, achieving an average accuracy of 89%. The framework was applied to over 6,000 enzyme sequences from the RemeDB to classify enzymes targeting diverse plastics, including PET, PLA, and Nylon. Besides, generative learning strategies combined with trained classification models in this work were applied for *de novo* generation of PET-degrading enzymes. Structural bioinformatics validated potential candidates through *in-silico* analysis, highlighting differences in physicochemical properties between generated and experimentally validated enzymes. Moreover, generated sequences exhibited lower molecular weights and higher aliphatic indices, features that may enhance interactions with hydrophobic plastic substrates. These findings highlight the utility of AI-based approaches in enzyme discovery, providing a scalable and efficient tool for addressing plastic pollution. Future work will focus on experimental validation of promising candidates and further refinement of generative strategies to optimize enzymatic performance.

## 1 Introduction

Plastic contamination poses a severe threat to the environment, impacting habitats, species, and ecosystems on a global scale (Li et al., 2021). This pollution occurs as plastics enter natural environments, endangering organisms, ecosystems, and even human health (MacLeod et al., 2021).

Plastics are synthetic polymers primarily derived from petrochemicals. They can be classified into two main categories: thermoplastics, which can be melted and reshaped multiple times (e.g., Polyethylene (PE), Polyethylene Terephthalate (PET), and Polypropylene (PP)), and thermosets, which after the curing process, cannot be reprocessed (e.g., Polyurethane (PU) and epoxy resins) (Bî rcă et al., 2019; Jadaun et al., 2022). The widespread use and persistence of plastics have significantly contributed to the accumulation of plastic waste, highlighting the urgent need for innovative and scalable solutions (Kumar et al., 2021).

Efforts to address plastic pollution include bioplastics, circular economy initiatives, and biodegradation (Folino et al., 2020). Bioplastics are derived from natural sources and offer eco-friendly alternatives but remain cost-prohibitive compared to traditional fossil-based plastics (Nandakumar et al., 2021). Circular economy practices, such as recycling and waste-to-energy conversion, have shown promise but are insufficient to mitigate the growing volume of plastic waste (Bhandari et al., 2021; Idumah and Chukwujike Nwuzor, 2019). Among these strategies, biodegradation emerges as a sustainable approach, leveraging the enzymatic activity of microorganisms to degrade plastics in an environmentally friendly manner (Folino et al., 2020).

Microbial enzymes capable of breaking down synthetic polymers have been identified, with hydrolases playing a critical role in plastic degradation (Yogalakshmi and Singh, 2019; Urbanek et al., 2020). However, advancing the discovery of plastic-degrading enzymes demands robust tools that not only identify enzyme candidates but also classify their substrate specificity. Recent advancements in bioinformatics and machine learning (ML) have begun to address these challenges, using techniques like metagenomic sequencing, protein language models, and structural predictions to explore enzyme functionality (Jiang et al., 2021; Jumper et al., 2021).

The development of machine learning-based classification models has significantly advanced the identification of plastic-degrading enzymes. Frameworks such as the Plastic Enzymatic Degradation (PED) model (Jiang et al., 2023) and PEZy-Miner (Jiang et al., 2024) integrate experimental data and computational tools to identify plastic-degrading enzymes. These models use protein sequence patterns and numerical representation techniques to achieve high prediction accuracy (Gupta and Agrawal, 2022). Additionally, machine learning-guided directed evolution (Wittmann et al., 2021) has optimized enzyme efficiency, such as enhancing PETase thermostability for improved PET degradation (Gupta and Agrawal, 2022; Lu et al., 2022). These approaches have accelerated enzyme discovery and enzyme design, providing computational solutions to explore extensive protein databases and uncover novel enzymes for biocatalytic plastic recycling.

Predictive models for plastic-degrading enzymes and their substrate detection are critical in addressing the complexities of plastic bioremediation. Given the vast diversity of enzyme sequences and plastic types, conventional methods such as cultivation or omics-based discovery are labor-intensive and time-consuming (Liu et al., 2021). Computational approaches, particularly machine learning models, fill this gap by offering scalable, efficient alternatives (Lu et al., 2022). These models not only enable the identification of enzymes with potential plastic-degrading capabilities but also predict substrate specificity, a key step in tailoring biocatalytic strategies to specific types of plastic waste (Taniguchi et al., 2019). The ability to classify enzyme-plastic interactions with high accuracy ensures a more targeted approach to mitigating plastic pollution and supports the development of engineered enzymes for industrial applications (Jiang et al., 2024).

This work introduces an integrated framework designed to facilitate the discovery and classification of plastic-degrading enzymes. The framework leverages fine-tuning methods with pre-trained protein language models to develop classification systems capable of detecting enzymes specific to seven types of plastics, including Nylon, PET, PLA, PCL, PHA, PHB, and PU/PUR. With an average accuracy of 89%, it demonstrates high robustness, making it a valuable tool for efficient enzyme functional annotation tools. The proposed framework successfully classified over 6,000 plastic-degrading enzymes from the RemeDB database (Sankara Subramanian et al., 2020). Additionally, generative learning strategies were incorporated for the *de novo* design of enzymes targeting specific plastic substrates, guided by Enzyme Commission (EC) numbers.

By combining structural bioinformatics and statistical analyses, the implemented framework evaluates both classified and generated sequences, positioning it as a powerful tool for accelerating the discovery and engineering of enzymes tailored to efficiently mitigate plastic pollution.

## 2 Methods

Diverse strategies and methodologies to advance the understanding and design of plastic-degrading enzymes were explored in this work. Curated datasets on enzyme-plastic interactions were generated through advanced data retrieval and processing techniques. Machine learning (ML) methods were then applied to develop binary classification models capable of identifying the specific plastic substrates targeted by these enzymes. To demonstrate the utility of the trained models, previously uncharacterized enzymes from the RemeDB database of Pollutant Degrading Enzymes from metagenomic sequences (Sankara Subramanian et al., 2020) were analyzed to predict their plastic substrates. Additionally, generative learning strategies, combined with the ML classification models, were employed to accelerate the *de novo* design of plastic-degrading enzymes and predict their substrate specificity. Finally, structural bioinformatics strategies were integrated to perform *in silico* validation of potential PET-degrading enzymes.

### 2.1 Collecting, preprocessing, and generating datasets

Data on plastic-degrading enzymes were collected using a keyword-based search strategy in Google Scholar and scientific repositories. Keywords such as “plastic-degrading enzymes”, “PET hydrolases”, and “biodegradation enzymes” were employed to identify relevant publications and databases. This approach enabled the retrieval of both experimental data and previously published datasets.

After identifying relevant sources, the datasets were downloaded, including experimental results, available datasets, and curated public databases like PlasticDB (Gambarini et al., 2022) and PAZy (Buchholz et al., 2022).

To ensure the datasets’ quality and compatibility with downstream analyses, several preprocessing steps were applied, including i) **Sequence extraction and annotation**: Only enzyme sequences with explicit annotations related to plastic degradation were retained, while non-annotated and duplicate entries were removed, ii) **Sequence length filtering**: Sequences shorter than 50 amino acids or longer than 1024 amino acids were excluded to maintain compatibility with numerical representation approaches, iii) **Inconsistency removal**: Sequences with conflicting annotations (e.g., labelled as both plastic-degrading and non-degrading) were excluded, and iv) **Non-canonical residues**: Sequences containing non-canonical amino acids were filtered out to facilitate the use of pre-trained protein language models.

Plastic types with fewer than 30 associated enzyme sequences were discarded to ensure sufficient data for classification model development (Biswas et al., 2021). The remaining data were consolidated into a single and integrated pivoted dataset linking enzyme sequences to their respective plastic-degrading activities. Additionally, supplementary enzyme sequences lacking specific plastic-type annotations were retrieved from RemeDB (Sankara Subramanian et al., 2020) and subjected to the same filtering and validation steps for consistency. These curated and validated datasets were subsequently used as input for training and testing classification models to predict the types of plastics they can degrade.

### 2.2 Machine learning strategies explored to develop binary classification models

This work investigates diverse strategies for developing binary classification models for detecting plastic substrates targeted by degrading enzymes, including i) traditional approaches that integrate supervised learning algorithms with feature engineering, ii) supervised learning algorithms combined with protein sequence encoding techniques, and iii) transfer learning strategies using pre-trained models as feature extraction methods alongside supervised learning algorithms.

The datasets for training the classification models were derived from the built pivoted dataset. Positive examples included enzyme-plastic substrate interactions. Negative examples included non-plastic-degrading enzymes reported in (Jiang et al., 2023) and randomly selected plastic-degrading enzymes without affinity for the chosen plastic substrate. To ensure balanced datasets, undersampling strategies were applied by randomly selecting negative examples. Finally, separate datasets were created for each plastic substrate analyzed in this work.

The pipeline for constructing classification models comprises i) a numerical representation strategy, ii) splitting the dataset into training and validation sets with a 75:25 ratio, iii) training the models using 10-fold cross-validation to mitigate overfitting, iv) evaluating performance metrics, and v) assessing model performance on the validation dataset.

Multiple methodologies were employed for numerical representation. Feature engineering strategies involved the use of the iFeature tool (Chen et al., 2018) alongside the MaxAbsoluteScaler for standardization and the SelectKBest method from the Scikit-learn library (Pedregosa et al., 2011) for feature selection. Encoding strategies included baseline methods such as one-hot encoding, frequency encoding, *k*-mer-based approaches, physicochemical property-based methods (Medina-Ortiz et al., 2022), and Fourier transform-based techniques (Siedhoff et al., 2021). Additionally, embeddings generated by pre-trained protein language models, such as ESM (Rives et al., 2021) and BERT (Kenton and Toutanova, 2019), were utilized as feature extraction methods.

The processed datasets were divided into training and validation/testing subsets. Various supervised learning algorithms were evaluated, exploring diverse hyperparameter configurations, including distance-based, rule-based, transformation-based, and ensemble-based methods. Cross-validation was employed to minimize overfitting, and model performance was evaluated using standard classification metrics such as accuracy, precision, recall, and F1-score (Section S1 of the Supplementary Materials for further details).

The training process was repeated 30 times to generate performance distributions and assess the stability of the models across different dataset partitions (Medina-Ortiz et al., 2024a). Models with superior average performance and minimal metric dispersion, indicating higher stability, were shortlisted for further evaluation (Medina-Ortiz et al., 2024b). These models were then tested on the validation dataset, and their performance metrics were documented.

After evaluating various strategies, the best-performing model was selected based on its consistent performance across training iterations, superior classification metrics, and minimal overfitting (Medina-Ortiz et al., 2024b).

### 2.3 Integrating trained binary classification models to discover plastic-degrading enzymes

The trained classification models were utilized to demonstrate their effectiveness in evaluating unknown plastic-degrading enzyme sequences and their integration into *de novo* generation workflows. Enzyme sequences were preprocessed using numerical representation, encoding, or descriptive techniques before applying the classification models. The models generated binary predictions, classifying each sequence as either positive or negative for the evaluated plastic substrate.

These models were applied to assess enzyme sequences obtained from the RemeDB database (Sankara Subramanian et al., 2020). The classification models were employed to evaluate the sequences and generate predictions.

To further integrate the classification models into *de novo* enzyme design. The ZymCTRL model (Munsamy et al., 2022) was used to generate enzyme sequences corresponding to seven selected EC numbers: 3.1.1.1, 3.1.1.2, 3.1.1.74, 3.1.1.102, 3.5.2.12, 3.5.1.46, and 3.5.1.117. For each EC number, over 15,000 sequences were generated. These *de novo* sequences were then processed and evaluated using the trained classification models to identify potential plastic-degrading enzymes and the plastic-substrate target. Finally, structural bioinformatics strategies were incorporated to perform *in silico* validation of the generated sequences.

### 2.4 Structural bioinformatic validation strategies for discovered PET enzymes through molecular docking methods

Structural bioinformatics analysis for PET-degrading enzymes was performed through molecular docking. As a baseline comparison, well-known structures of experimentally validated PET-degrading enzymes—PETase (PDB ID: 6ANE) and DuraPETase (PDB ID: 6KY5)—were used. Additionally, proteins reported as plastic-degrading enzymes in the RemeDB database with known structures (Sankara Subramanian et al., 2020), and identified by the classification models trained in this study as having the highest probability of PET-degradation activity, were included. These enzyme structures were retrieved from the Protein Data Bank (PDB) using their respective UniProt accession codes. Moreover, *de novo* generated sequences—whose structures were unknown—were also analyzed. The *de novo* sequences predicted to be PET-degrading enzymes by the classification model with the highest probability were selected for structural bioinformatic validation. Additionally, generated enzymes lacking PET-degrading activity were included. These structures were predicted using ColabFold v1.5.5 (Mirdita et al., 2022).

As a ligand, two polyethylene terephthalate (2PET) molecules were used as input and prepared using the AutoDock Tools v1.5.6 software (Goodsell and Olson, 1990) to generate the required pdbqt file.

All protein structures were preprocessed using PyMOL 3.1 (Yuan et al., 2017) by applying i) isolating of protein chains, ii) removing water molecules, and iii) centering the structures. Then, the AutoDock prepare receptor4 module (Morris et al., 2009) was employed to add hydrogens, protonate the structures, and convert them to the pdbqt format required for docking.

Different docking grids were defined depending on the input protein structure. For PETase, and DuraPETase, the docking grid was defined around binding sites previously reported in (Fecker et al., 2018; Cui et al., 2019). In PETase, residues Y60, T61, A62, W132, S133, M134, W158, D179, I181, H210, and S211 were considered, while for DuraPETase, the binding site was defined by residues Y87, F117, Y119, W185, and H214. In contrast, for proteins obtained from RemeDB and the generated enzymes, binding sites were predicted using Fpocket v3.0 (Le Guilloux et al., 2009).

Molecular docking was processed using VINA v1.2.0 software (Forli et al., 2016). The resulting docking poses were exported in pdb format and clustered based on the root-mean-square deviation (RMSD). Finally, the optimal docking pose was identified by considering multiple factors, including the most populated cluster, the highest number of interactions with the binding site residues, and the lowest docking score.

### 2.5 Implementation strategies

The source code was developed in Python (version 3.12). Classification models using supervised learning algorithms were constructed with the scikit-learn library (Pedregosa et al., 2011). Integration of pre-trained protein language models was implemented using the Transformers library (Wolf et al., 2020) and the bio-embedding library (Dallago et al., 2021). Generative models were based on the pre-trained ZymCTRL model (Munsamy et al., 2022). A dedicated Conda environment was created to facilitate the deployment and reproducibility of the work. Moreover, various Jupyter notebooks were developed to facilitate and demonstrate how to use the models and how to integrate on *de-novo* enzyme design strategies or exploration of unknown enzymes.

## 3 Results and Discussion

### 3.1 More than 400 plastic-degrading enzymes with known plastic-substrate targets were identified

An integrated dataset comprising over 400 experimentally validated plastic-degrading enzymes with known target plastic-type was generated by compiling information from public databases, including PlasticDB (Gambarini et al., 2022), PAZy (Buchholz et al., 2022), and PMDB (Gan and Zhang, 2019). Additionally, public datasets were incorporated to generate the integrated dataset (Jiang et al., 2023; Gambarini et al., 2021; Tournier et al., 2023). The built dataset includes over 30 types of plastic substrates, such as PET, PHB, PHA, and Nylon. Among these, enzymes capable of degrading PET and PHB are the most represented, with 183 and 70 enzymes, respectively (Table 1).

**Table 1:**
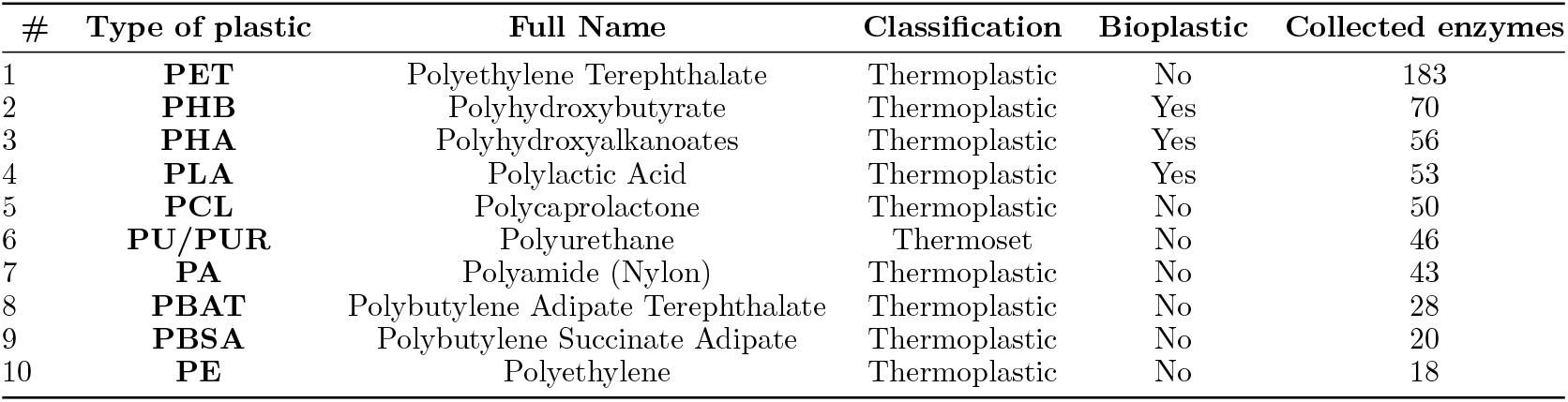
Summary of the top ten plastic-substrate targets identified in the integrated dataset. The full name, classification as thermoset or thermoplastic, biodegradability status, and the number of associated collected examples are included.

Table 1 provides an overview of the top ten plastic types represented in the integrated dataset, highlighting key properties such as their classification (thermoplastic or thermoset), biodegradability, and the number of associated enzymes collected for each type. Polyethylene Terephthalate (PET) stands out as the most represented plastic, with 183 enzymes capable of degrading it. Bioplastics such as Polyhydroxybutyrate (PHB), Polyhydroxyalkanoates (PHA), and Polylactic Acid (PLA) are also well-represented in the dataset, with 70, 56, and 53 enzymes, respectively, indicating a growing interest in biodegradable alternatives to conventional plastics.

The integrated dataset demonstrates the dominance of thermoplastics, which are more readily recycled and enzymatically degraded than chemically stable thermosets (Podara et al., 2024; Oladele et al., 2023). Only polyurethane (PU/PUR) is represented in thermosets, with 46 associated enzymes. Inherently non-biodegradable plastics such as Polyethylene (PE) and Polyamide (Nylon) are also included, though with fewer associated enzymes (Bergeson et al., 2024; Ghatge et al., 2020).

Less common plastics are also presented, broadening the dataset’s scope. Polybutylene Succinate (PBS), a biodegradable thermoplastic, leads this category with 16 enzymes. Other bioplastics like Polyhydroxybutyrate-co-Valerate (PHBV), Polyvinyl Alcohol (PVA), and Polyethylene Furanoate (PEF) are present with 7, 5, and 4 enzymes, respectively.

The dataset illustrates remarkable promiscuity by identifying some enzymes that have shown activity for degrading multiple plastic types. Sixty-five enzymes target exactly two types of plastics, 19 degrade three types, and 27 can act on more than three. The most frequent two-plastic combinations are PHA and PHB, with 44 enzymes reflecting their structural similarity and shared biodegradability. Other notable pairs include PCL with PET or PLA, each involving 22 enzymes. For three-plastic combinations, the most common involves PCL, PHA, and PLA, with 15 enzymes, followed by combinations such as PCL with PES and PHA, each supported by 11 enzymes. Niche combinations, such as ECOFLEX, PBSA, and PLA or PHA, PHBH, and PHBVH, occur less frequently, with only one enzyme identified for each. The plastic degradation capacities of the enzymes examined are illustrated in Figure 1, which shows the main combinations of plastics each enzyme can degrade and provides an overview of their prevalence within the dataset. In addition, Supplementary Table S1 provides a detailed overview of enzyme degradation capabilities for each combination of plastic types.

**Figure 1.**
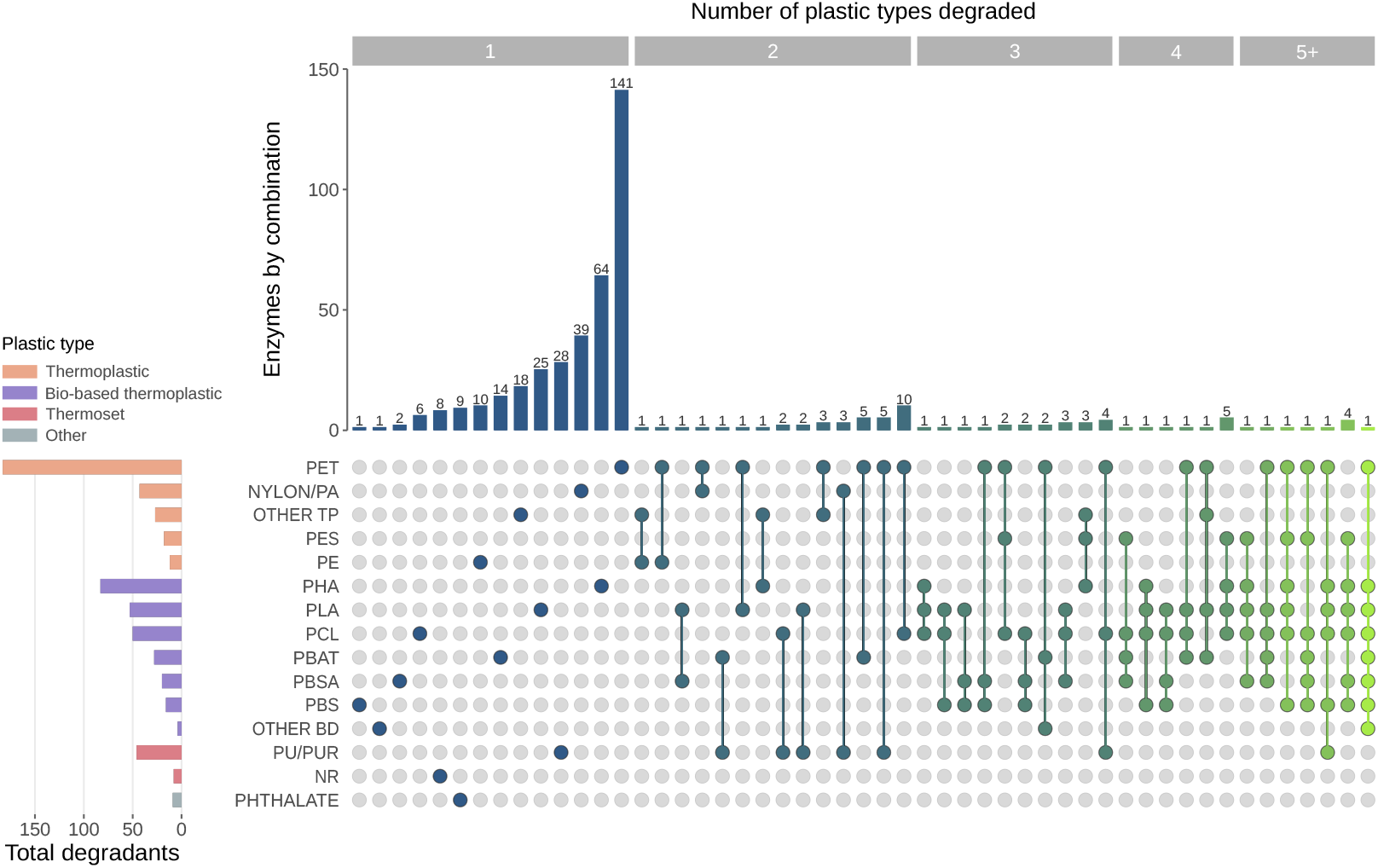
Enzyme distribution based on their plastic degradation capabilities. The matrix illustrates the specific combinations of plastic types that the evaluated enzymes can degrade, using colored dots to indicate degradation. Above the matrix, a histogram reflects the number of enzymes linked to each combination, with colors representing their promiscuity levels. The bar chart on the left shows the total number of degraded plastic types, with bars coloured according to plastic classification. Plastics with few observations were categorized as either “OTHER TP” (thermoplastics) or “OTHER BD” (bio-based thermoplastics), while PHA variants were consolidated into the “PHA” category. Of the enzymes studied, 82.6% degrade only one plastic-type, while the rest show a range of promiscuity, with the ability to degrade up to eight different plastic types.

### 3.2 Seven binary classification models were trained for detecting plastic-substrate target of degrading enzymes with performance over 85% of precision

Binary classification models were developed to identify the plastic substrates targeted by plastic-degrading enzymes. Using the generated dataset and the positive example length filter outlined in the methodology, seven plastic-substrate targets were selected for training, including Nylon, PCL, PET, PHA, PHB, PLA, and PU/PUR (Table 2).

**Table 2:** Performance of selected model for each classification model for identifying the plastic substrates targeted by plastic-degrading enzymes.

| Type of plastic | Algorithm | Embedding | F1-score | Accuracy | Precision | Recall | MCC | Sensitivity | Specificity |
| --- | --- | --- | --- | --- | --- | --- | --- | --- | --- |
| PET | ExtraTrees ensemble | Protrans XLU50 | 0.94+/- 0.02 | 0.94 +/- 0.02 | 0.95 +/- 0.03 | 0.94+/- 0.04 | 0.89+/- 0.04 | 0.94 +/- 0.04 | 0.95 +/- 0.03 |
| PHB | ExtraTrees ensemble | ESM-1b | 0.91 +/- 0.06 | 0.93 +/- 0.04 | 0.93 +/- 0.08 | 0.91 +/- 0.08 | 0.86 +/- 0.09 | 0.91 +/- 0.08 | 0.95 +/- 0.05 |
| PHA | Bagging | ESM-e | 0.90 +/- 0.06 | 0.90 +/- 0.06 | 0.92 +/- 0.05 | 0.87 +/- 0.07 | 0.79 +/- 0.12 | 0.87 +/- 0.07 | 0.92 +/- 0.06 |
| PLA | Hist Gradient Boosting | Plusrnn | 0.81 +/- 0.07 | 0.81 +/- 0.07 | 0.76 +/- 0.10 | 0.87 +/- 0.10 | 0.64 +/- 0.13 | 0.87 +/- 0.10 | 0.75 +/- 0.11 |
| PCL | Random Forest | Protrans BDF | 0.84 +/- 0.06 | 0.83+/- 0.06 | 0.82 +/- 0.11 | 0.88 +/- 0.08 | 0.66 +/- 0.13 | 0.88 +/- 0.08 | 0.78 +/- 0.13 |
| PU/PUR | Random Forest | Protrans AIBERT | 0.83 +/- 0.05 | 0.82 +/- 0.05 | 0.77 +/- 0.10 | 0.92 +/- 0.05 | 0.66 +/- 0.10 | 0.92 +/- 0.05 | 0.74 +/- 0.14 |
| PA (Nylon) | Random Forest | ESM-e | 0.97 +/- 0.02 | 0.97 +/- 0.02 | 0.98 +/- 0.03 | 0.96 +/- 0.04 | 0.94 +/- 0.05 | 0.96 +/- 0.04 | 0.99 +/- 0.03 |

During the exploration phase, over 7,000 binary classification models were evaluated for each plastic type by combining various numerical representation strategies with supervised learning algorithms (Section S2 of Supplementary Materials for details). Embedding-based strategies consistently outperformed baseline methods such as one-hot encoding, feature-engineering approaches, and physicochemical-based encoders, demonstrating the superior utility of embeddings for functional classification tasks.

Within embedding-based strategies, models trained with embeddings derived from pre-trained protein language models like ESM (Rives et al., 2021) and ProTrans (Elnaggar et al., 2021) achieved the best performance, with average F1-scores exceeding 0.8 in both training and validation stages. In contrast, embeddings generated from pre-trained models, like CPCProt (Lu et al., 2020), GloVe (Pennington et al., 2014), and FastText (Joulin et al., 2016), showed lower performance, with F1-scores below 0.75.

The performance of embedding-based models also varied depending on the supervised learning algorithms used (Section S2 of Supplementary Materials for more details). Ensemble-based algorithms like Random Forest and Adaboost achieved superior results compared to Linear Discriminant Analysis and Quadratic Discriminant Analysis. Aligning with the trends in embedding performance, the top algorithms achieved average F1-scores above 0.8, whereas lower-performing models failed to exceed an F1-score of 0.75.

When selecting the best-performing strategies, specific combinations of pre-trained models and supervised learning algorithms were identified for different plastic-substrate targets. For Nylon, embeddings derived from ESM and ProTrans, combined with algorithms like Extra-Trees and Random Forest, achieved the highest performance, with average MCC values of 0.94 and 0.92, respectively. For PU/PUR, the best results were obtained with ProTrans embeddings paired with Random Forest and ESM embeddings combined with Random Forest (Section S2 of Supplementary Materials for more details).

To identify the best models for each plastic type, selection criteria focused on maximizing performance, ensuring stability across training partitions, and minimizing overfitting were applied (Medina-Ortiz et al., 2024a,b). These criteria prioritized models that demonstrated high accuracy and strong generalization capabilities. Table 2 summarizes the selected models, detailing the ML algorithm, representation strategy, and corresponding performance metrics for each plastic-substrate target.

PET and PA achieve the highest performance with Extra-Trees ensembles combined with ProTrans XLU50 and Random Forest combined with ESM-e embeddings, respectively. Both models achieve F1-scores and accuracy exceeding 0.94, with low standard deviations, reflecting stability and strong generalization.

For PHB and PHA, Extra-Trees and Bagging algorithms with ESM-based embeddings deliver solid results, with F1-scores around 0.90. However, slightly lower MCC values indicate reduced consistency compared to PET and PA models. Models for PLA (Hist Gradient Boosting with PlusRNN embeddings), PCL (Random Forest with ProTrans BDF), and PU/PUR (Random Forest with ProTrans AlBERT embeddings) show moderate performance, with F1-scores of 0.81, 0.84, and 0.83, respectively, and higher variability in metrics such as MCC and specificity.

Figure 2 summarizes the performance evaluation of the trained models, showing variation across different plastic-substrate targets. For high-performing models like PET, PA/NYLON, and PHA, both the ROC and Precision-Recall (PR) curves demonstrate the best results. These models exhibit strong discrimination capabilities with minimal false positives or false negatives, as evident in the confusion matrices.

**Figure 2.**
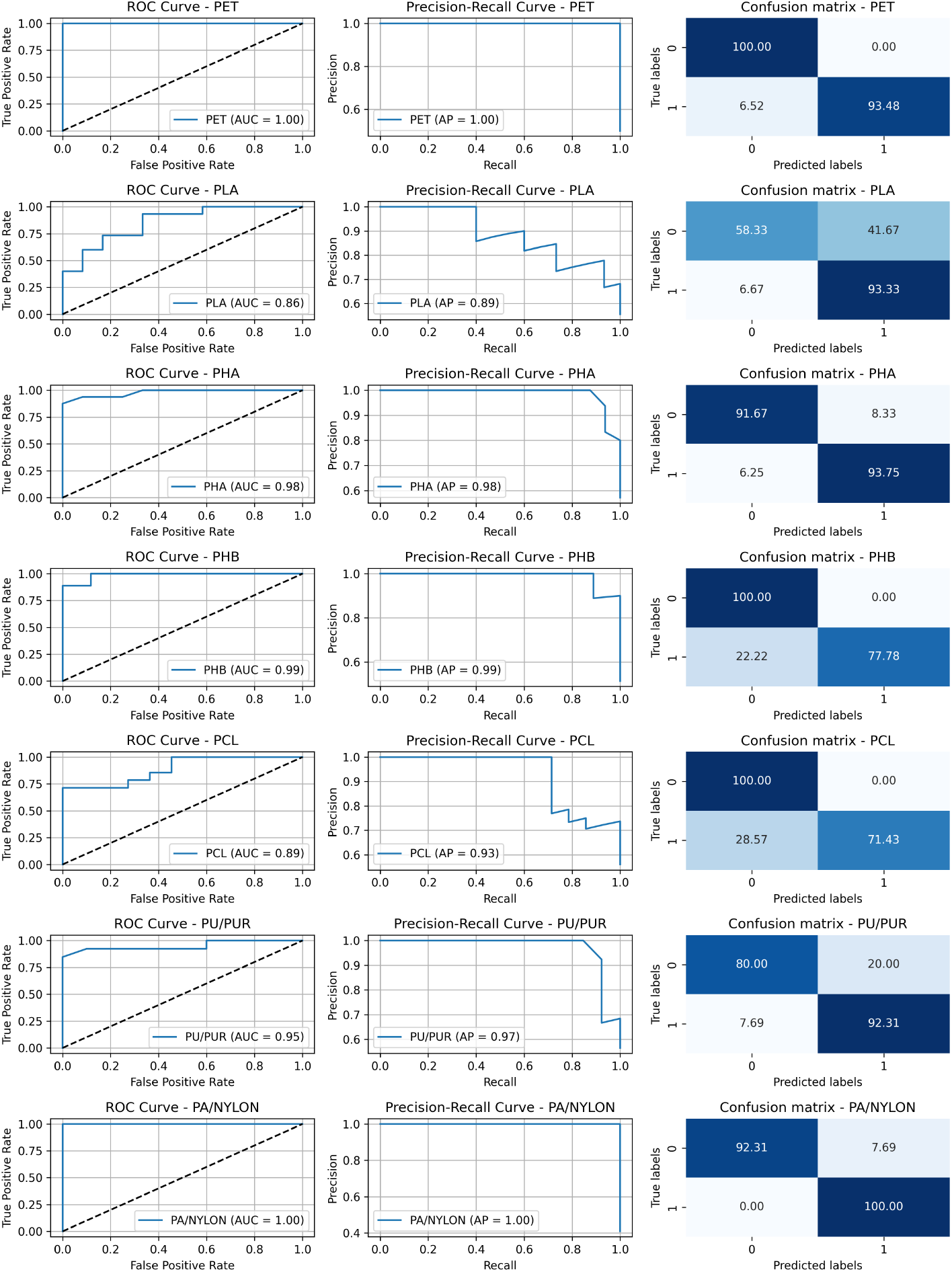
Performance description of seven binary classification models for identifying the plastic-substrate targets by plastic-degrading enzymes. The trained models were evaluated using ROC curves, precision-recall curves, and confusion matrices. Among the models, those designed to detect PET, PA, and PHA demonstrated high robustness, with high AUC and average precision scores, as well as minimal misclassifications. Conversely, the models for PHB and PCL exhibited higher false negative rates, indicating a tendency to miss true positive samples. Similarly, the models for PLA and PU/PUR displayed elevated false positive rates, suggesting challenges in discriminating between true negatives and false positives.

On the other hand, models like PLA, PHB, and PCL, while still achieving good performance, exhibit more variability. The AUC for these models remains high (e.g., 0.86 for PLA, 0.99 for PHB, and 0.89 for PCL), but their confusion matrices reveal limitations. PLA shows a higher proportion of false positives, with 41.67% of incorrect classifications. In contrast, PCL and PHB show higher false negative rates, indicating that a noticeable portion of true samples was not identified.

### 3.3 Evaluating unknown plastic-degrading enzymes collected from RemeDB

More than 6,000 plastic-degrading enzymes were collected and processed from the RemeDB database (Sankara Subramanian et al., 2020) to demonstrate the application of the trained models in identifying plastic-substrate targets. Initially, the enzyme sequences were converted into numerical representations using embeddings extracted from pre-trained protein language models. These embeddings served as inputs for the trained classification models, which generated probability scores for both the positive and negative classes. A default threshold (probability *>*0.5) was then applied to classify the enzymes, allowing for the identification of their potential plastic-substrate targets. Alternatively, different thresholds were also applied to explore how to change the identified enzymes for each plastic-substrate target explored in this work.

Table 3 presents the number of enzymes identified for different plastic-substrate targets when evaluating unknown sequences from RemeDB (Sankara Subramanian et al., 2020) across various probability thresholds. A closer analysis reveals that models like PLA and PU/PUR, which showed high false positive rates during validation, tend to identify a larger number of sequences, particularly at lower thresholds. For example, the PLA model identifies 2,133 sequences at a 50% threshold and still retains 626 sequences at the stricter 90% threshold. The elevated false positive rates observed during validation suggest that these predictions could include significant errors, reducing the reliability of the identified sequences.

**Table 3:**
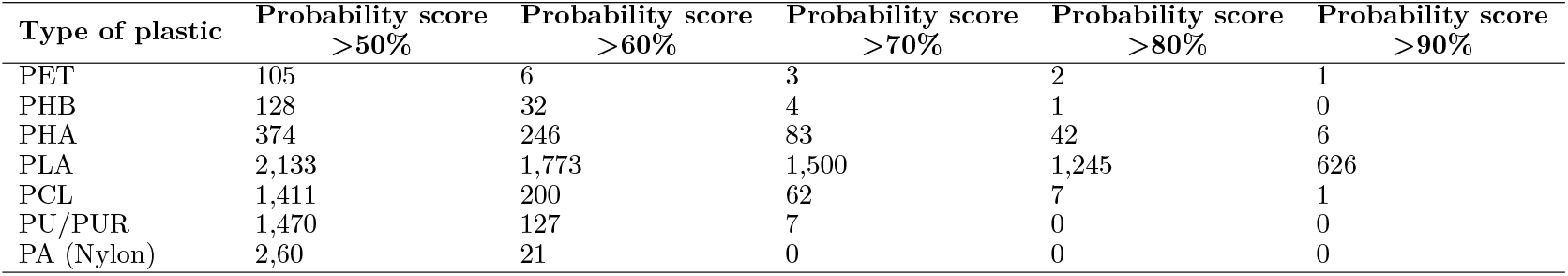
Identified enzymes for each plastic-substrate target evaluated in this work for the collected enzymes from RemeDB (Sankara Subramanian et al., 2020)

| Type of plastic | Probability score<br>>50% | Probability score<br>>60% | Probability score<br>>70% | Probability score<br>>80% | Probability score<br>>90% |
| --- | --- | --- | --- | --- | --- |
| PET | 105 | 6 | 3 | 2 | 1 |
| PHB | 128 | 32 | 4 | 1 | 0 |
| PHA | 374 | 246 | 83 | 42 | 6 |
| PLA | 2,133 | 1,773 | 1,500 | 1,245 | 626 |
| PCL | 1,411 | 200 | 62 | 7 | 1 |
| PU/PUR | 1,470 | 127 | 7 | 0 | 0 |
| PA (Nylon) | 2,60 | 21 | 0 | 0 | 0 |

In contrast, models like PET, PHB, and PCL, which exhibited lower false positive rates, demonstrate a more conservative identification of sequences. The PET model identifies only 105 sequences at a 50% threshold, and this number drops to a single sequence at a 90% threshold. While this may appear as a limitation, it indicates that the sequences identified at higher thresholds are likely more reliable, consistent with the validation results showing low error rates.

### 3.4 Integrating generative learning for *de novo* design of PET-degrading enzymes with *in silico* validation

This work explores the integration of generative learning strategies with trained classification models to identify plastic-substrate targets for plastic-degrading enzymes. The combined approach demonstrates the potential of generative models to expand the landscape of enzyme candidates and enhance the discovery process.

Using the pre-trained model implemented by Munsamy et al. (2022), alongside the EC number classification described in the methodology, more than 100,000 unique enzyme sequences were generated. These sequences were subsequently processed using pre-trained protein language models to extract embeddings, followed by the application of the PET classification model to identify potential PET-degrading enzymes. As a result, over 2,900 enzymes were detected as PET-degrading candidates.

A comparative analysis of amino acid frequencies between the generated and experimentally validated PET-degrading enzymes derived from the dataset constructed in this work (Section S3 of Supplementary Materials) reveals interesting trends. While the majority of residues showed minimal differences, alanine and glycine were observed in higher proportions in the generated sequences, whereas serine was underrepresented compared to the validated sequences.

Further analysis of physicochemical properties highlights clear differences between the generated and experimentally validated sequences (Section S3 of Supplementary Materials). The experimentally validated sequences exhibited lower values for molecular weight, isoelectric point, charge density, net charge, instability index, Boman index, and aromaticity. These differences suggest that the generated enzymes may possess reduced structural complexity and charge-related characteristics. In contrast, higher aliphatic index and hydrophobic ratio values were observed in the generated sequences, indicating a preference for aliphatic residues and greater hydrophobicity. These traits could influence the functionality and stability of the generated enzymes, potentially enhancing interactions with hydrophobic substrates like PET.

The differences observed suggest that generative strategies produce enzymes with distinct physicochemical profiles compared to experimentally validated enzymes. While these differences could represent novel functional variations, they also highlight the need for experimental validation to determine the catalytic efficiency, stability, and substrate specificity of these generated candidates.

Structural bioinformatics strategies were employed for the *in silico* validation of explored PET-degrading enzymes, encompassing both the potential PET-degrading enzymes identified in RemeDB (Sankara Subramanian et al., 2020) and the *de novo* sequences generated through generative learning approaches. Molecular dynamics analyses were conducted for experimentally validated PET-degrading enzymes (PETase and DuraPETase), the most promising candidates identified by the classification model, and generated enzymes lacking PET-degrading activity, as summarized in Figure 3.

**Figure 3.**
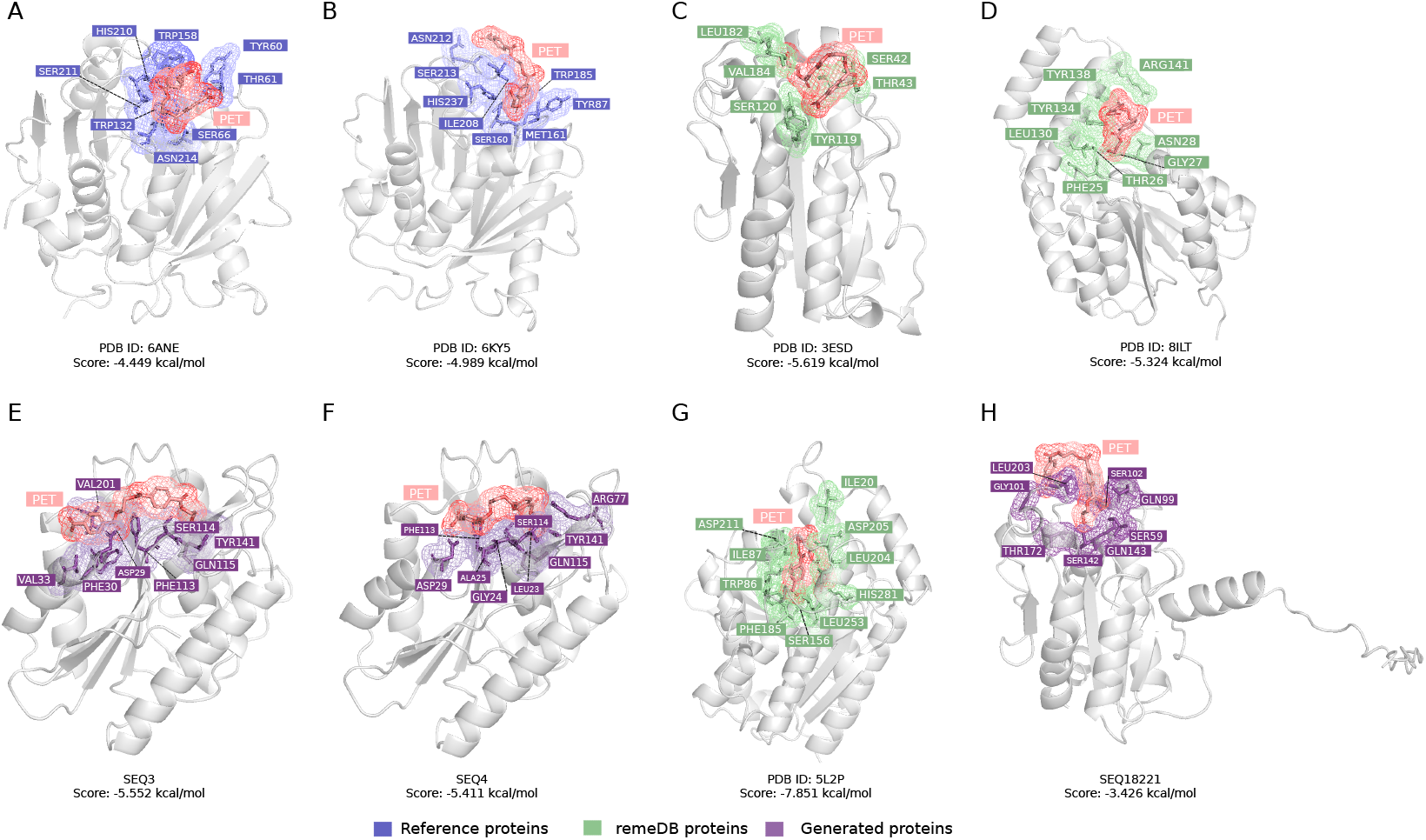
Structural analysis through molecular dynamics approach to compare interactions between PET substrate and plastic-degrading enzymes. **(A-B)** Best docking poses and key residues for experimentally validated PET-degrading enzymes, PETase (PDB ID: 6ANE) and DuraPETase (PDB ID: 6KY5), respectively, highlighting their established catalytic activity and substrate interactions. **(C-D)** Best docking poses and key residues for potential PET-degrading enzymes identified from the RemeDB database (Sankara Subramanian et al., 2020), as predicted using the trained classification models. **(E-F)** Best docking poses and key residues for potential PET-degrading enzymes generated through the pre-trained generative model of Munsamy et al. (2022), showcasing novel enzyme-substrate interactions derived from artificial enzyme design.**(G)** A promising PET-degrading enzyme identified from RemeDB (Sankara Subramanian et al., 2020) with the lowest docking score, indicating the highest affinity between the PET substrate and enzyme. **(H)** A generated enzyme, predicted as non-PET-degrading, with a high docking score suggesting a low affinity for the PET substrate, underscoring its limited degradation potential.

The results obtained through docking reveal interactions between the studied enzymes and the PET substrate, highlighting frequent residues in the binding sites such as SER, TYR, and ASP. For PETase and DuraPETase enzymes (Figure 3 (A-B)), the interaction patterns observed were consistent with those reported in the literature. For PETase, residues such as TYR60, TYR61, TRP132, TRP158, and HIS210 were identified. Specifically, TYR60 and TRP158 have been reported to form an aromatic clamp that stabilizes the terephthalic ring within the active site, facilitating its interaction in a highly folded conformation (Fecker et al., 2018).

Similarly, for the potential PET-degrading enzymes identified from RemeDB (Sankara Subramanian et al., 2020), interactions involving tyrosine residues in the active site were also observed, stabilizing the interaction with the substrate (Figure 3 (C-D)).

On the other hand, for the generated sequences identified as PET degrading enzymes by the classification model (Figure 3 (E-F)), differences are observed in the conformation of the PET substrate within the site. In case E, PET adopts an extended pose, stabilized by potential hydrophobic interactions between VAL201 and VAL33 with the aliphatic chains of the substrate, while the aromatic residues PHE30, PHE113, and TYR141 establish *π − π* interactions with the rings. These interactions are complemented by possible hydrogen bonds with ASP29, SER114, and GLN115. In contrast, in case F, PET exhibits an intermediate conformation, neither fully extended nor highly folded. Here, the aromatic residues PHE113 and TYR141 maintain *π − π* interactions with the ring, while ASP29, GLN115, and ARG77 contribute through hydrogen bonds and possible electrostatic interactions.

An analysis of extreme cases was also conducted to explore the limits of the model predictions. Figure 3 G illustrates the best docking pose for a potential PET-degrading enzyme identified from RemeDB (Sankara Subramanian et al., 2020). This enzyme exhibited the lowest docking score of -7.851 kcal/mol, indicating the highest binding affinity between the enzyme and the PET substrate.

Additionally, a negative case was evaluated using enzymes generated by the pre-trained generative model. Figure 3 H shows the best docking pose for a non-PET-degrading enzyme classified by the PET-classification model, which had a low probability value for PET degradation (0.32). This docking pose demonstrated the highest docking score (indicating the lowest binding affinity), highlighting its reduced potential for substrate interaction.

## 4 Conclusions

This work presents a comprehensive AI-driven framework combining machine learning and generative modeling to accelerate the discovery of plastic-degrading enzymes. A curated dataset was constructed by integrating data from PlasticDB, PAZy, PMDB databases, and other public datasets of plastic-degrading enzymes, collecting more than 400 sequences of experimentally validated enzymes targeting 30 different types of plastics. The plastics with the most reported enzymes are PET, PHB, and PHA.

Binary classification models were trained for seven different plastic substrates (PET, PHB, PHA, PLA, PCL, PU, and PA) using the integrated dataset of plastic-degrading enzymes. The framework demonstrated high performance and robustness, achieving an accuracy ranging from 81% to 97% across the different plastic targets. The developed models successfully classified over 6,000 plastic-degrading enzymes from sequences deposited in the RemeDB. Furthermore, the analysis was expanded by integrating generative learning to *de novo* design over 100,000 enzyme candidates. Among them, 2,900 were identified as potential PET-degrading enzymes according to the trained classification model.

Analysis of physicochemical properties revealed distinct differences between generated and experimentally validated enzymes, such as lower molecular weight, reduced aromaticity, and higher aliphatic index in generated sequences. These variations suggest potential functional advantages for interacting with hydrophobic plastics like PET. *In-silico* validation confirmed the structural feasibility of selected candidates, offering insights into their degradation potential.

The present work provides a valuable tool for discovering new plastic-degrading enzymes, facilitating and accelerating wet lab efforts in the characterization of putative plastic-degrading enzymes by scoring and categorizing potential candidates.

Future work will focus on experimental validation of promising enzyme candidates to confirm their catalytic activity and stability. Additionally, advanced generative techniques, coupled with iterative optimization and experimental feedback, will be explored to refine enzyme properties further. This integrative approach represents a significant step toward scalable and sustainable solutions for mitigating plastic pollution through biocatalysis.

## Data availability statement

The source code, environment configuration, datasets, and trained models are freely available for non-commercial use in the GitHub repository at https://github.com/ProteinEngineering-PESB2/PetDegradation under the MIT license.

## Author contributions statement

DM-O, AD and DA-S: conceptualization. DM-O, NS-G, and DA-S: methodology. DM-O, AD, BA, and JA-A: validation. DM-O, DA-S, NS, JA and AD: investigation. DM-O, DA-S, NS-G, JA, SR, DS-V, BA, JA-A, and AD: writing, review, and editing. BA and JA-A: supervision and funding resources. BA and JA-A: project administration.

## Acknowledgments

DM-O acknowledges ANID for the project “SUBVENCIÓN A INSTALACIÓN EN LA ACADEMIA CONVOCATORIA AÑO 2022”, Folio 85220004. AD acknowledges ANID for funding through the “FONDECYT Iniciación project 11230208”. DS-V acknowledges ANID for the scholarship “Beca doctorado nacional convocatoria año 2018”, Folio 21180560. SR thanks ANID for the Doctorate Fellowship (21221519). DM-O, AD, SR, DS-V, BA, and JA-A gratefully acknowledge support from ANID (Exploración project 13220166) and the Centre for Biotechnology and Bioengineering - CeBiB (PIA project FB0001, AFB240001, ANID, Chile).

## References

Bergeson, A. R., Silvera, A. J., and Alper, H. S. (2024). Bottlenecks in biobased approaches to plastic degradation. Nature Communications, 15(1):4715.

Bhandari, N. L., Bhattarai, S., Bhandari, G., Subedi, S., and Dhakal, K. N. (2021). A Review on Current Practices of Plastics Waste Management and Future Prospects. Journal of Institute of Science and Technology, 26(1):107–118.

Biswas, S., Khimulya, G., Alley, E. C., Esvelt, K. M., and Church, G. M. (2021). Low-n protein engineering with data-efficient deep learning. Nature methods, 18(4):389–396.

Buchholz, P. C., Feuerriegel, G., Zhang, H., Perez-Garcia, P., Nover, L.-L., Chow, J., Streit, W. R., and Pleiss, J. (2022). Plastics degradation by hydrolytic enzymes: The plastics-active enzymes database—pazy. Proteins: Structure, Function, and Bioinformatics, 90(7):1443–1456.

Bîrča, A., Gherasim, O., Grumezescu, V., and Grumezescu, A. M. (2019). Chapter 1 - introduction in thermoplastic and thermosetting polymers. In Grumezescu, V. and Grumezescu, A. M., editors, Materials for Biomedical Engineering, pages 1–28. Elsevier.

Chen, Z., Zhao, P., Li, F., Leier, A., Marquez-Lago, T. T., Wang, Y., Webb, G. I., Smith, A. I., Daly, R. J., Chou, K.-C., et al. (2018). ifeature: a python package and web server for features extraction and selection from protein and peptide sequences. Bioinformatics, 34(14):2499–2502.

Cui, Y., Chen, Y., Liu, X., Dong, S., Tian, Y., Qiao, Y., Mitra, R., Han, J., Li, C., Han, X., et al. (2019). Computational redesign of a petase for plastic biodegradation by the grape strategy. BioRxiv, page 787069.

Dallago, C., Schütze, K., Heinzinger, M., Olenyi, T., Littmann, M., Lu, A. X., Yang, K. K., Min, S., Yoon, S., Morton, J. T., and Rost, B. (2021). Learned embeddings from deep learning to visualize and predict protein sets. Current Protocols, 1(5):e113.

Elnaggar, A., Heinzinger, M., Dallago, C., Rehawi, G., Wang, Y., Jones, L., Gibbs, T., Feher, T., Angerer, C., Steinegger, M., Bhowmik, D., and Rost, B. (2021). Prottrans: Towards cracking the language of life’s code through self-supervised deep learning and high performance computing. IEEE Transactions on Pattern Analysis and Machine Intelligence, 44(10):7112–7127.

Fecker, T., Galaz-Davison, P., Engelberger, F., Narui, Y., Sotomayor, M., Parra, L. P., and Ramírez-Sarmiento, C. A. (2018). Active site flexibility as a hallmark for efficient pet degradation by i. sakaiensis petase. Biophysical journal, 114(6):1302–1312.

Folino, A., Karageorgiou, A., Calabro, P. S., and Komilis, D. (2020). Biodegradation of wasted bioplastics in natural and industrial environments: A review. Sustainability, 12(15).

Forli, S., Huey, R., Pique, M. E., Sanner, M. F., Goodsell, D. S., and Olson, A. J. (2016). Computational protein–ligand docking and virtual drug screening with the autodock suite. Nature protocols, 11(5):905– 919.

Gambarini, V., Pantos, O., Kingsbury, J. M., Weaver, L., Handley, K. M., and Lear, G. (2021). Phylogenetic distribution of plastic-degrading microorganisms. Msystems, 6(1):10–1128.

Gambarini, V., Pantos, O., Kingsbury, J. M., Weaver, L., Handley, K. M., and Lear, G. (2022). Plasticdb: a database of microorganisms and proteins linked to plastic biodegradation. Database, 2022:baac008.

Gan, Z. and Zhang, H. (2019). Pmbd: a comprehensive plastics microbial biodegradation database. Database, 2019:baz119.

Ghatge, S., Yang, Y., Ahn, J.-H., and Hur, H.-G. (2020). Biodegradation of polyethylene: a brief review. Applied Biological Chemistry, 63:1–14.

Goodsell, D. S. and Olson, A. J. (1990). Automated docking of substrates to proteins by simulated annealing. Proteins: Structure, Function, and Bioinformatics, 8(3):195–202.

Gupta, A. and Agrawal, S. (2022). Machine Learning-Based Enzyme Engineering of PETase for Improved Efficiency in Degrading Non-Biodegradable Plastic. bioRxiv.

Idumah, C. and Chukwujike Nwuzor, I. (2019). Novel trends in plastic waste management. SN Applied Sciences, 1.

Jadaun, J. S., Bansal, S., Sonthalia, A., Rai, A. K., and Singh, S. P. (2022). Biodegradation of plastics for sustainable environment. Bioresource Technology, 347:126697.

Jiang, R., Shang, L., Wang, R., Wang, D., and Wei, N. (2023). Machine learning based prediction of enzymatic degradation of plastics using encoded protein sequence and effective feature representation. Environmental Science & Technology Letters, 10(7):557–564.

Jiang, R., Yue, Z., Shang, L., Wang, D., and Wei, N. (2024). Pezy-miner: An artificial intelligence driven approach for the discovery of plastic-degrading enzyme candidates. Metabolic Engineering Communications, page e00248.

Jiang, Y. et al. (2021). Machine learning methods for enzyme function prediction. Bioinformatics Advances.

Joulin, A., Grave, E., Bojanowski, P., and Mikolov, T. (2016). Bag of tricks for efficient text classification. arXiv preprint 1607.01759.

Jumper, J., Evans, R., Pritzel, A., Green, T., Figurnov, M., Ronneberger, O., Tunyasuvunakool, K., Bates, R., Žídek, A., Potapenko, A., et al. (2021). Highly accurate protein structure prediction with alphafold. Nature, 596(7873):583–589.

Kenton, J. D. M.-W. C. and Toutanova, L. K. (2019). Bert: Pre-training of deep bidirectional transformers for language understanding. In Proceedings of naacL-HLT, volume 1, page 2. Minneapolis, Minnesota.

Kumar, R., Verma, A., Shome, A., Sinha, R., Sinha, S., Jha, P., Kumar, R., Kumar, P., Shubham Das, S., Sharma, P., and Prasad, P. V. V. (2021). Impacts of plastic pollution on ecosystem services, sustainable development goals, and need to focus on circular economy and policy interventions. Sustainability.

Le Guilloux, V., Schmidtke, P., and Tuffery, P. (2009). Fpocket: an open source platform for ligand pocket detection. BMC bioinformatics, 10:1–11.

Li, P., Wang, X., Su, M., Zou, X., Duan, L., and Zhang, H. (2021). Characteristics of plastic pollution in the environment: A review. Bulletin of Environmental Contamination and Toxicology, 107.

Liu, Y. et al. (2021). Machine learning approaches in petase enzyme discovery and engineering. Biotechnology Advances, 49:107752.

Lu, A. X., Zhang, H., Ghassemi, M., and Moses, A. (2020). Self-supervised contrastive learning of protein representations by mutual information maximization. BioRxiv, pages 2020–09.

Lu, H., Diaz, D. J., Czarnecki, N. J., Zhu, C., Kim, W., Shroff, R., Acosta, D. J., Alexander, B. R., Cole, H. O., Zhang, Y., et al. (2022). Machine learning-aided engineering of hydrolases for pet depolymerization. Nature, 604(7907):662–667.

MacLeod, M., Arp, H. P., Tekman, M. B., and Jahnke, A. (2021). The global threat from plastic pollution. Science, 373:61.

Medina-Ortiz, D., Cabas-Mora, G., Moya-Barria, I., Soto-Garcia, N., and Uribe-Paredes, R. (2024a). Rudeus, a machine learning classification system to study dna-binding proteins. bioRxiv, pages 2024–02.

Medina-Ortiz, D., Contreras, S., Amado-Hinojosa, J., Torres-Almonacid, J., Asenjo, J. A., Navarrete, M., and Olivera-Nappa, Á. (2022). Generalized property-based encoders and digital signal processing facilitate predictive tasks in protein engineering. Frontiers in Molecular Biosciences, 9.

Medina-Ortiz, D., Contreras, S., Fernández, D., Soto-García, N., Moya, I., Cabas-Mora, G., and Olivera-Nappa, Á. (2024b). Protein language models and machine learning facilitate the identification of antimicrobial peptides. International Journal of Molecular Sciences, 25(16):8851.

Mirdita, M., Schütze, K., Moriwaki, Y., Heo, L., Ovchinnikov, S., and Steinegger, M. (2022). Colabfold: making protein folding accessible to all. Nature methods, 19(6):679–682.

Morris, G. M., Huey, R., Lindstrom, W., Sanner, M. F., Belew, R. K., Goodsell, D. S., and Olson, A. J. (2009). Autodock4 and autodocktools4: Automated docking with selective receptor flexibility. Journal of computational chemistry, 30(16):2785–2791.

Munsamy, G., Lindner, S., Lorenz, P., and Ferruz, N. (2022). Zymctrl: a conditional language model for the controllable generation of artificial enzymes. In NeurIPS Machine Learning in Structural Biology Workshop.

Nandakumar, A., Chuah, J.-A., and Sudesh, K. (2021). Bioplastics: A boon or bane? Renewable and Sustainable Energy Reviews, 147:111237.

Oladele, I. O., Okoro, C. J., Taiwo, A. S., Onuh, L. N., Agbeboh, N. I., Balogun, O. P., Olubambi, P. A., and Lephuthing, S. S. (2023). Modern trends in recycling waste thermoplastics and their prospective applications: a review. Journal of Composites Science, 7(5):198.

Pedregosa, F., Varoquaux, G., Gramfort, A., Michel, V., Thirion, B., Grisel, O., Blondel, M., Prettenhofer, P., Weiss, R., Dubourg, V., et al. (2011). Scikit-learn: Machine learning in python. the Journal of machine Learning research, 12:2825–2830.

Pennington, J., Socher, R., and Manning, C. D. (2014). Glove: Global vectors for word representation. In Proceedings of the 2014 conference on empirical methods in natural language processing (EMNLP), pages 1532–1543.

Podara, C., Termine, S., Modestou, M., Semitekolos, D., Tsirogiannis, C., Karamitrou, M., Trompeta, A.-F., Milickovic, T. K., and Charitidis, C. (2024). Recent trends of recycling and upcycling of polymers and composites: A comprehensive review. Recycling, 9(3):37.

Rives, A., Meier, J., Sercu, T., Goyal, S., Lin, Z., Guo, D., Ott, M., Zitnick, C. L., Ma, J., and Fergus, R. (2021). Biological structure and function emerge from scaling unsupervised learning to 250 million protein sequences. Proceedings of the National Academy of Sciences, 118(15).

Sankara Subramanian, S. H., Balachandran, K. R. S., Rangamaran, V. R., and Gopal, D. (2020). Remedb: tool for rapid prediction of enzymes involved in bioremediation from high-throughput metagenome data sets. Journal of Computational Biology, 27(7):1020–1029.

Siedhoff, N. E., Illig, A.-M., Schwaneberg, U., and Davari, M. D. (2021). Pypef—an integrated framework for data-driven protein engineering. Journal of Chemical Information and Modeling, 61(7):3463–3476.

Taniguchi, I., Yoshida, S., Hiraga, K., Miyamoto, K., Kimura, Y., and Oda, K. (2019). Biodegradation of PET: Current Status and Application Aspects. ACS Catalysis, 9(5):4089–4105.

Tournier, V., Duquesne, S., Guillamot, F., Cramail, H., Taton, D., Marty, A., and André, I. (2023). Enzymes’ power for plastics degradation. Chemical Reviews, 123(9):5612–5701.

Urbanek, A. K., Mironćzuk, A. M., García-Martín, A., Saborido, A., de la Mata, I., and Arroyo, M. (2020). Biochemical properties and biotechnological applications of microbial enzymes involved in the degradation of polyester-type plastics. Biochimica et Biophysica Acta (BBA) - Proteins and Proteomics, 1868(2):140315.

Wittmann, B. J., Johnston, K. E., Wu, Z., and Arnold, F. H. (2021). Advances in machine learning for directed evolution. Current opinion in structural biology, 69:11–18.

Wolf, T., Debut, L., Sanh, V., Chaumond, J., Delangue, C., Moi, A., Cistac, P., Rault, T., Louf, R., Funtowicz, M., Davison, J., Shleifer, S., von Platen, P., Ma, C., Jernite, Y., Plu, J., Xu, C., Scao, T. L., Gugger, S., Drame, M., Lhoest, Q., and Rush, A. M. (2020). Transformers: State-of-the-art natural language processing. In Proceedings of the 2020 Conference on Empirical Methods in Natural Language Processing: System Demonstrations, pages 38–45, Online. Association for Computational Linguistics.

Yogalakshmi, K. N. and Singh, S. (2019). Plastic Waste: Environmental Hazards, Its Biodegradation, and Challenges, pages 99–133. Springer Singapore.

Yuan, S., Chan, H. S., and Hu, Z. (2017). Using pymol as a platform for computational drug design. Wiley Interdisciplinary Reviews: Computational Molecular Science, 7(2):e1298.

