## Supplementary Materials for "Protein language models accelerate the discovery of Plastic-Degrading Enzymes"

#### S1 Metrics performances used to evaluate the classification models

Let $TP$ be the true positive values (correctly classified by the model), $FP$ be the false positive (unexpected results), the $FN$ the false negative (missing result), and the $TN$ are the true negative (correctly absent of result). Using the defined ratios, it is possible to determine the more common metrics employed to evaluate the performance of a predictive model. The equations 1, 2, 3, 4, and 5 represent the classic metrics employed to evaluate the performance of the predictive models for classification tasks and the metrics utilised in this work. Besides, sensitivity and specificity were also used in this work, estimated by employing the equations 6 and 7, respectively.

$$accuracy = \frac{TP + TN}{TP + FP + TN + FN} \quad (1)$$

$$precision = \frac{TP}{TP + FP} \quad (2)$$

$$recall = \frac{TP}{TP + FN} \quad (3)$$

$$F_\beta = \frac{(1+\beta^2)TP}{(1+\beta^2)TP + FP + \beta^2 FN} \quad (4)$$

$$MCC = \frac{(TP \times TN) - (FP \times FN)}{\sqrt{(TP + FP)(TP + FN)(TN + FP)(TN + FN)}} \quad (5)$$

$$Sensitivity = \frac{TP}{TP + FN} \quad (6)$$

$$Specificity = \frac{TN}{TN + FP} \quad (7)$$

*

#### S2 Enzyme degradation capabilities

Supplementary Table **S1** provides a comprehensive aggregation of enzymes characterized by the combinations of plastics they degrade, as analyzed within the dataset.

| Degradation promiscuity | Plastics | Enzymes |
|---|---|---|
| 1 | PET | 141 |
| | NYLON/PA | 39 |
| | PU/PUR | 28 |
| | PHB | 25 |
| | PLA | 25 |
| | PBAT | 14 |
| | PE | 10 |
| | PHTHALATE | 9 |
| | NR | 8 |
| | TP | 8 |
| | PCL | 6 |
| | PHA | 3 |
| | PVA | 3 |
| | LDPE | 2 |
| | PBSA | 2 |
| | PEG | 2 |
| | PBS | 1 |
| | PMCL | 1 |
| | PS | 1 |
| 2 | PHA, PHB | 28 |
| | PCL, PET | 10 |
| | PBAT, PET | 5 |
| | PET, PU/PUR | 5 |
| | PEF, PET | 3 |
| | PU/PUR, NYLON/PA | 3 |
| | O-PVA, PVA | 2 |
| | PCL, PU/PUR | 2 |
| | PLA, PU/PUR | 2 |
| | LDPE, PE | 1 |
| | PBAT, PU/PUR | 1 |
| | PBS-BLEND, PBSA-BLEND | 1 |
| | PBSA, PLA | 1 |
| | PE, PET | 1 |
| | PET, NYLON/PA | 1 |
| | PET, PLA | 1 |
| | PHA, PHO | 1 |
| | PHB, PHBV | 1 |
| 3 | PCL, PET, PU/PUR | 4 |
| | PBSA, PCL, PLA | 3 |
| | PBS, PBSA, PCL | 2 |
| | PCL, PES, PET | 2 |
| | PHA, PHB, PHV | 2 |
| | P3HV, PHA, PHBV | 1 |
| | PBS, PBSA, PET | 1 |
| | PBS, PBSA, PLA | 1 |
| | PBS, PCL, PLA | 1 |
| | PHA, PHB, PHBV | 1 |
| | PHA, PHB, PPL | 1 |
| 4 | ECOVIO-FT, PBAT, PBSET, PET | 2 |
| | PCL, PES, PHA, PLA | 2 |
| | P(3HB-CO-3MP), PHA, PHB, PHV | 1 |
| | PBAT, PBSA, PCL, PES | 1 |
| | PBAT, PCL, PET, PLA | 1 |
| | PBAT, PEF, PET, PLA | 1 |
| | PBS, PBSA, PCL, PLA | 1 |
| 5 | PBAT, PBSA, PCL, PET, PLA | 1 |
| | PBS, PCL, PHA, PHB, PLA | 1 |
| | PCL, PES, PHA, PHBV, PLA | 1 |
| 6 | P3HP, P4HB, PEA, PES, PHA, PHB | 3 |
| | PCL, PES, PHA, PHBV, PHO, PLA | 2 |
| | PBAT, PBS, PBSA, PCL, PES, PET | 1 |
| | PBS, PBSA, PCL, PES, PHA, PLA | 1 |
| | PBSA, PCL, PES, PHA, PHBV, PLA | 1 |
| | PCL, PHA, PHB, PHBH, PHBVH, PLA | 1 |
| 7 | PBS, PBSA, PCL, PES, PHA, PHB, PLA | 3 |
| | PBS, PCL, PES, PET, PHA, PHB, PLA | 1 |
| | PBS, PCL, PET, PHA, PHB, PLA, PU/PUR | 1 |
| 9 | ECOFLEX, PBAT, PBS, PBSA, PCL, PET, PHA, PHB, PLA | 1 |

**Supplementary Table S1**: Detailed summary of enzymes degradation capabilities grouped by combinations of plastic types.

### S3 Training binary classification models for plastic-substrate targets using machine learning strategies

#### S3.1 Classification models for PET

Figure **S1** summarizes the performances obtained during explorations strategies by type of encoder approaches. Figure **S2** summarizes the performances by encoder strategy. Figure **S3** summarizes the performances by supervised learning algorithms, and Figure **S4** summarizes the performances by type of pre-trained protein language models (embedding).

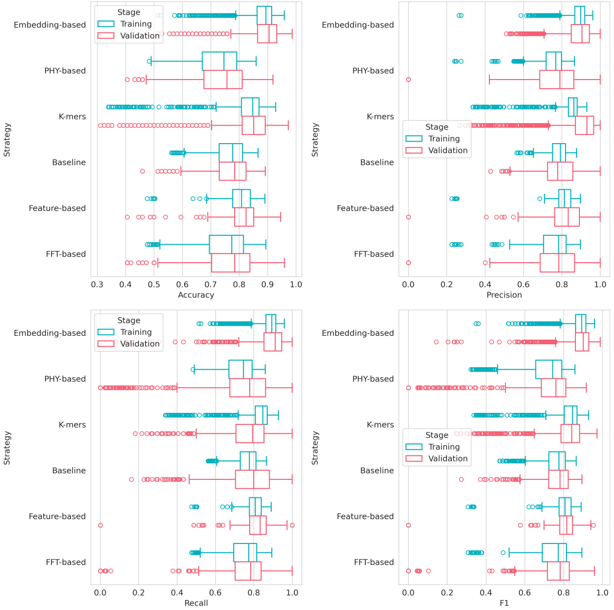


**Supplementary Figure S1**: Distribution performances for explored modes evaluated by type of encoder

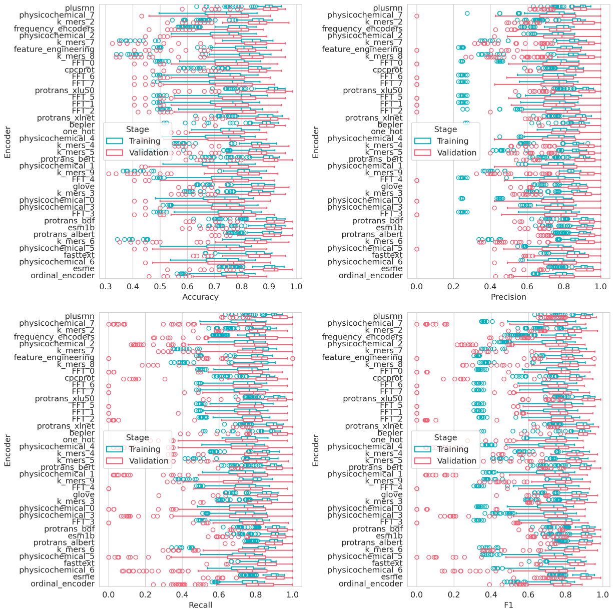


**Supplementary Figure S2**: Distribution performances for explored modes evaluated by encoder

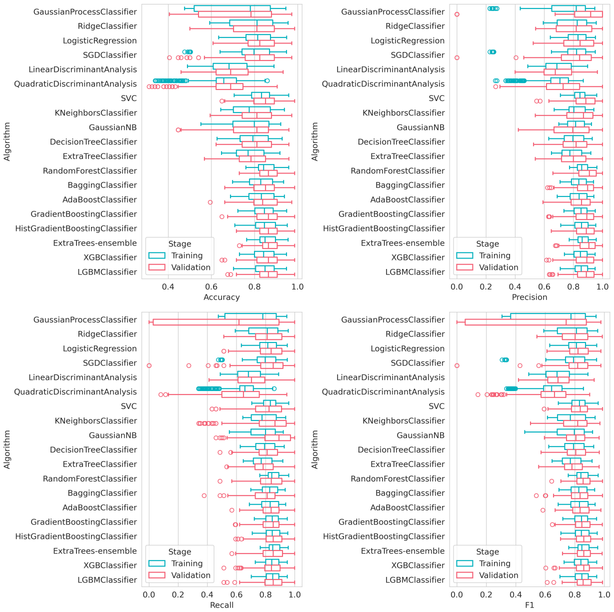


**Supplementary Figure S3**: Distribution performances for explored modes evaluated by algorithm

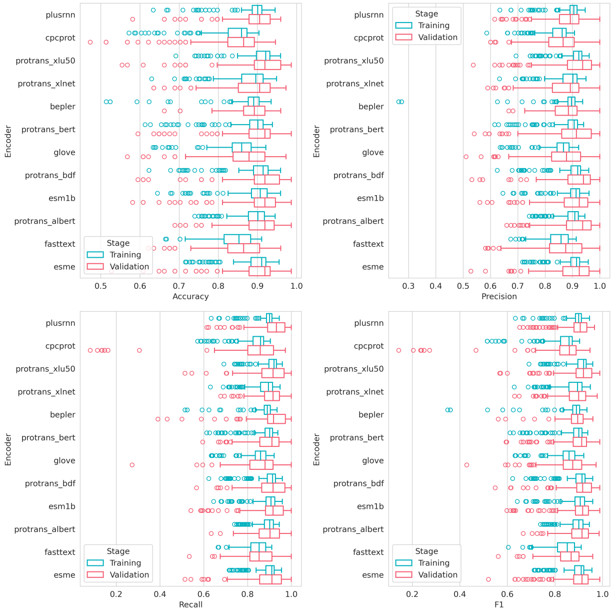


**Supplementary Figure S4**: Distribution performances for explored modes evaluated by type of pre-trained model

#### S3.2 Classification models for PLA

Figure **S5** summarizes the performances obtained during explorations strategies by type of encoder approaches. Figure **S6** summarizes the performances by encoder strategy. Figure **S7** summarizes the performances by supervised learning algorithms, and Figure **S8** summarizes the performances by type of pre-trained protein language models (embedding).

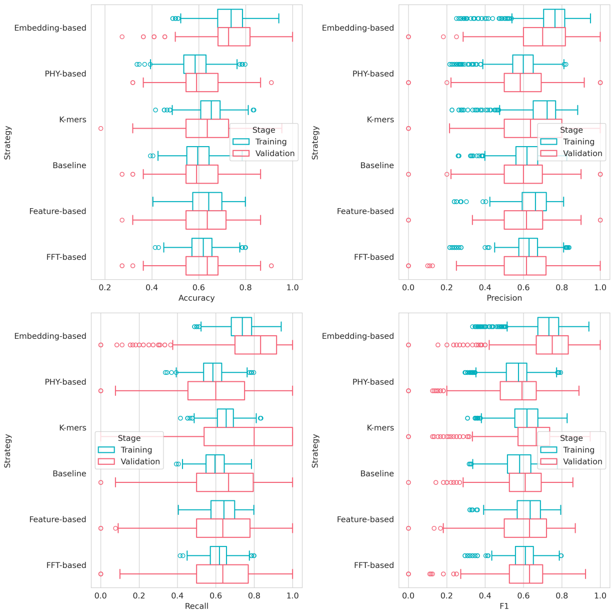


**Supplementary Figure S5**: Distribution performances for explored modes evaluated by type of encoder

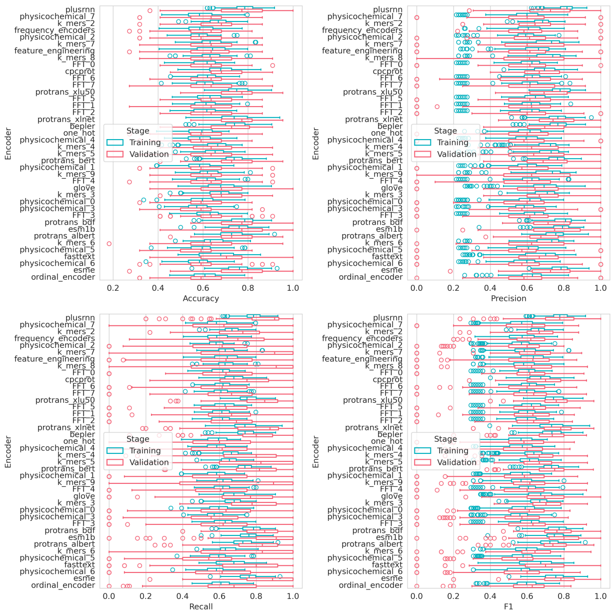


**Supplementary Figure S6**: Distribution performances for explored modes evaluated by encoder

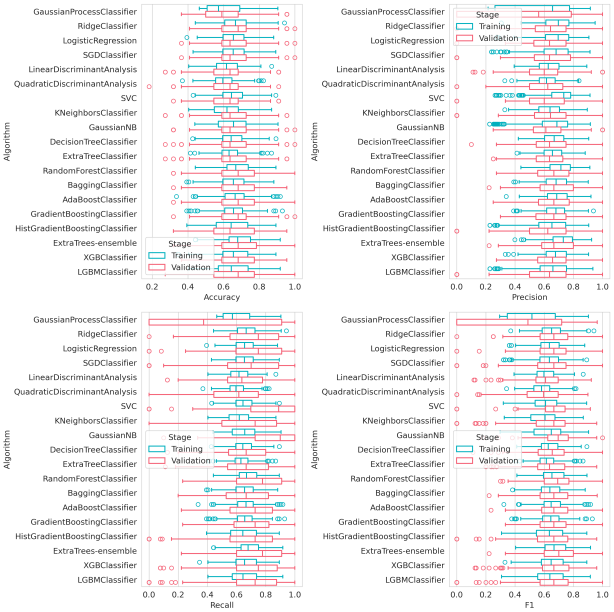


**Supplementary Figure S7**: Distribution performances for explored modes evaluated by algorithm

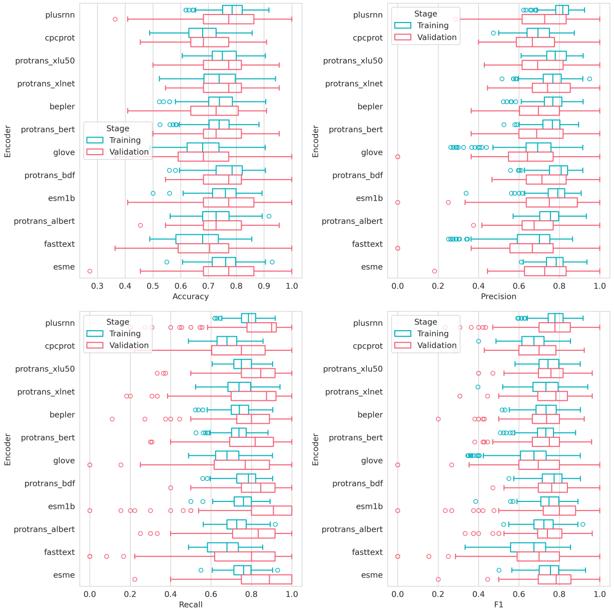


**Supplementary Figure S8**: Distribution performances for explored modes evaluated by type of pre-trained model

#### S3.3 Classification models for PHA

Figure **S9** summarizes the performances obtained during explorations strategies by type of encoder approaches. Figure **S10** summarizes the performances by encoder strategy. Figure **S11** summarizes the performances by supervised learning algorithms, and Figure **S12** summarizes the performances by type of pre-trained protein language models (embedding).

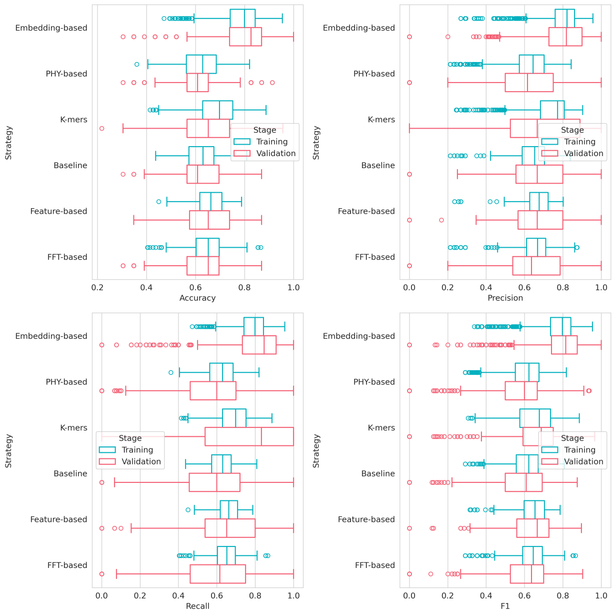


**Supplementary Figure S9**: Distribution performances for explored modes evaluated by type of encoder

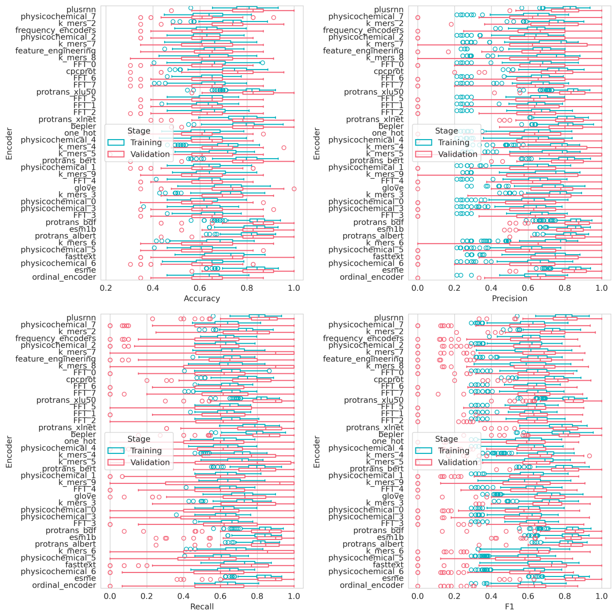


**Supplementary Figure S10**: Distribution performances for explored modes evaluated by encoder

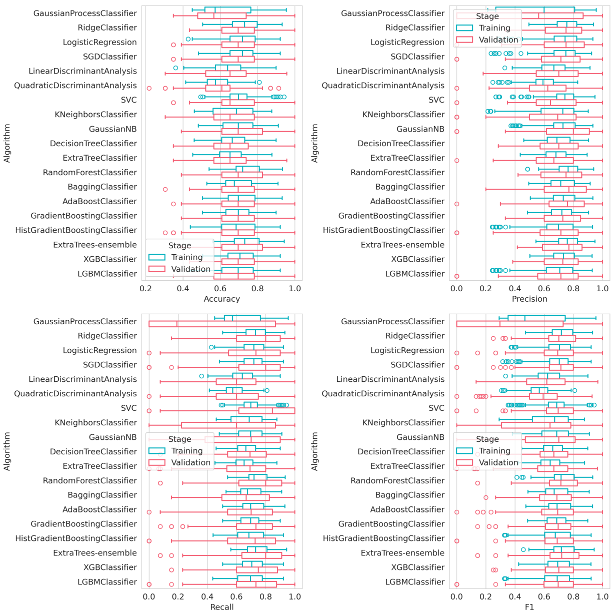


**Supplementary Figure S11**: Distribution performances for explored modes evaluated by algorithm

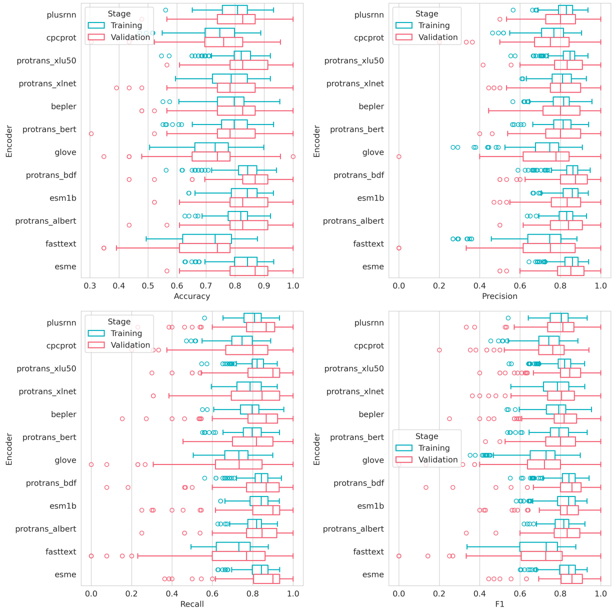


**Supplementary Figure S12**: Distribution performances for explored modes evaluated by type of pre-trained model

#### S3.4 Classification models for PHB

Figure **S13** summarizes the performances obtained during explorations strategies by type of encoder approaches. Figure **S14** summarizes the performances by encoder strategy. Figure **S15** summarizes the performances by supervised learning algorithms, and Figure **S16** summarizes the performances by type of pre-trained protein language models (embedding).

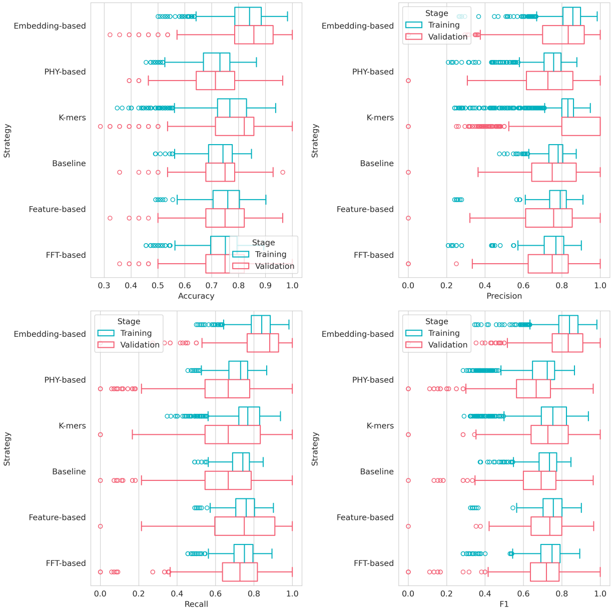


**Supplementary Figure S13**: Distribution performances for explored modes evaluated by type of encoder

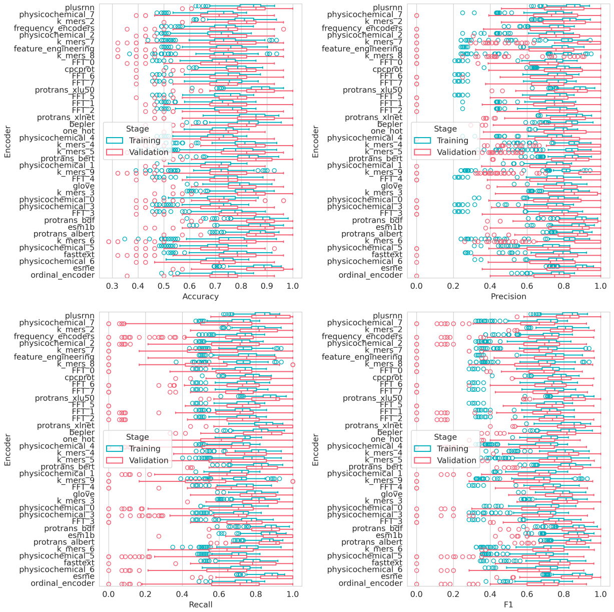


**Supplementary Figure S14**: Distribution performances for explored modes evaluated by encoder

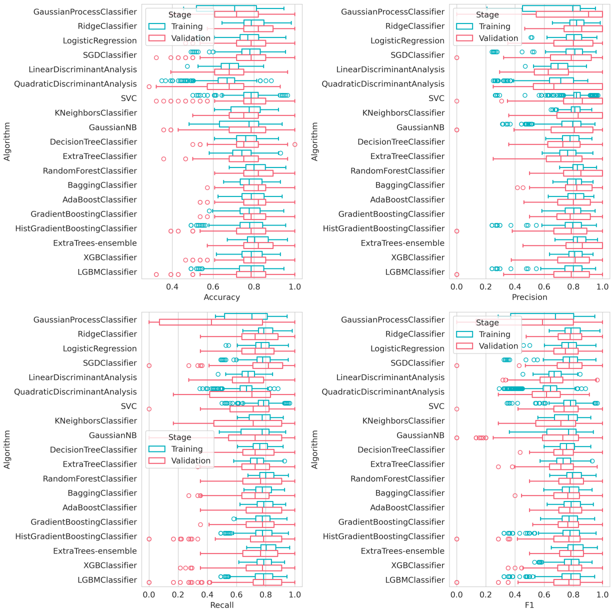


**Supplementary Figure S15**: Distribution performances for explored modes evaluated by algorithm

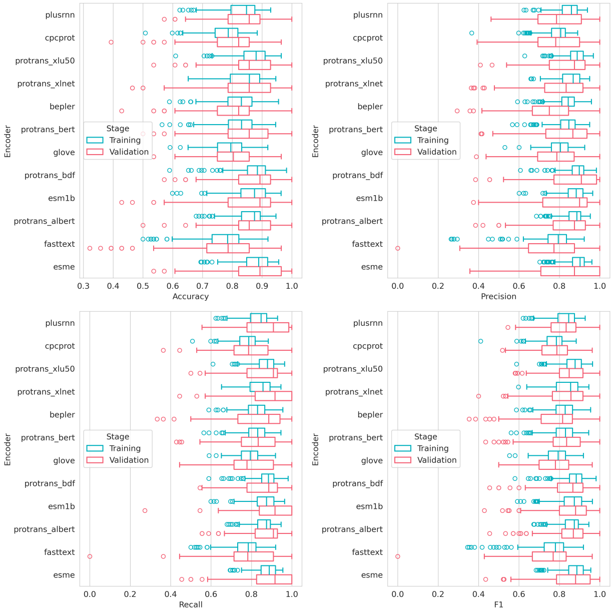


**Supplementary Figure S16**: Distribution performances for explored modes evaluated by type of pre-trained model

#### S3.5 Classification models for PCL

Figure **S17** summarizes the performances obtained during explorations strategies by type of encoder approaches. Figure **S18** summarizes the performances by encoder strategy. Figure **S19** summarizes the performances by supervised learning algorithms, and Figure **S20** summarizes the performances by type of pre-trained protein language models (embedding).

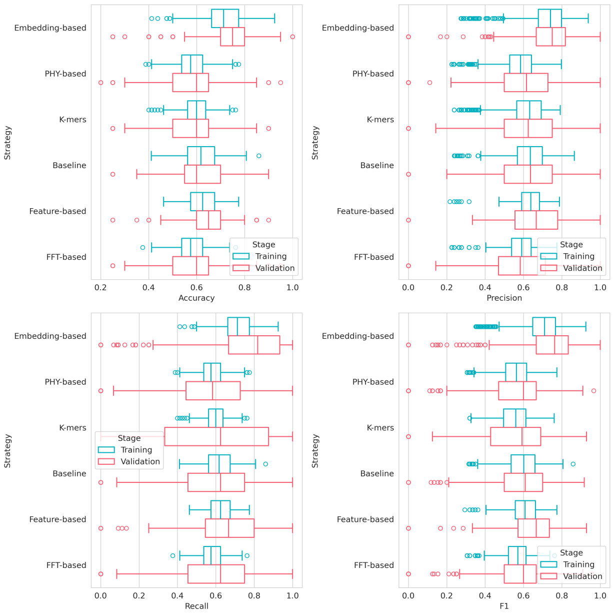


**Supplementary Figure S17**: Distribution performances for explored modes evaluated by type of encoder

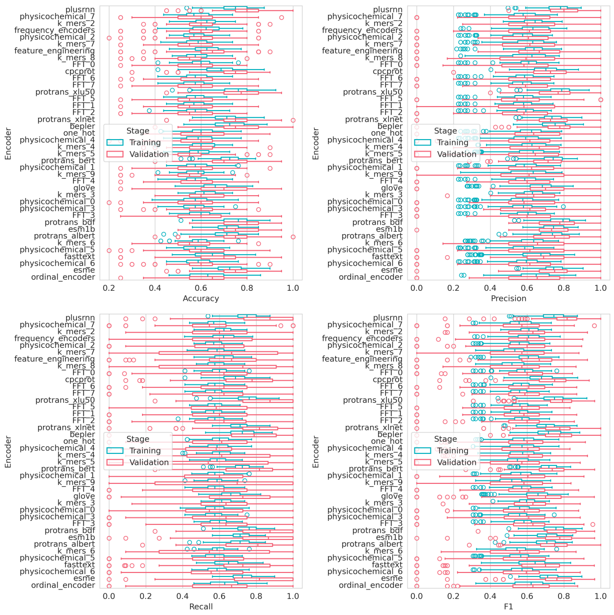


**Supplementary Figure S18**: Distribution performances for explored modes evaluated by encoder

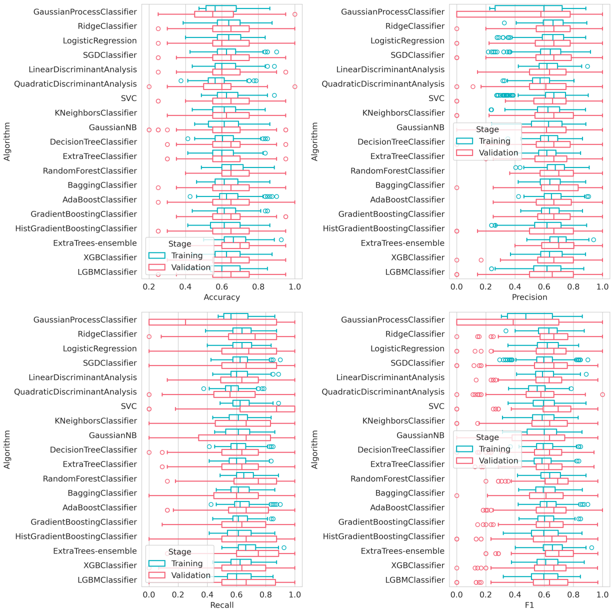


**Supplementary Figure S19**: Distribution performances for explored modes evaluated by algorithm

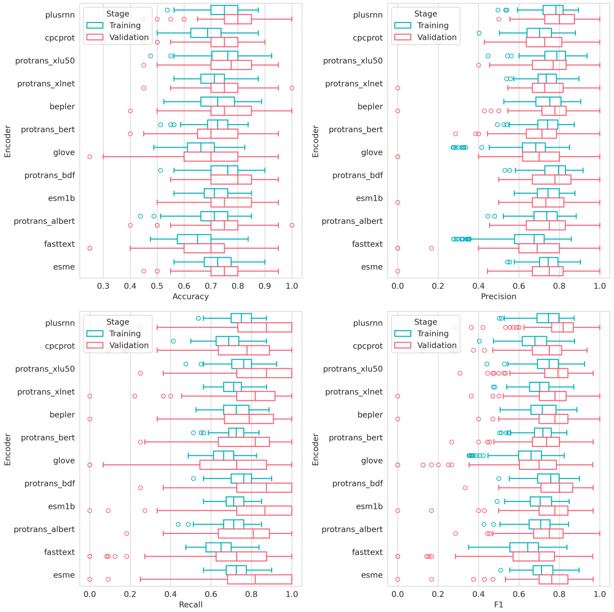


**Supplementary Figure S20**: Distribution performances for explored modes evaluated by type of pre-trained model

#### S3.6 Classification models for Nylon

Figure **S21** summarizes the performances obtained during explorations strategies by type of encoder approaches. Figure **S22** summarizes the performances by encoder strategy. Figure **S23** summarizes the performances by supervised learning algorithms, and Figure **S24** summarizes the performances by type of pre-trained protein language models (embedding).

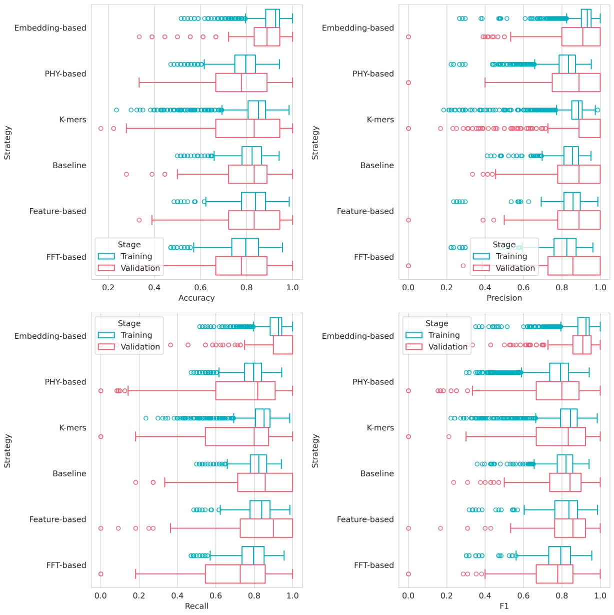


**Supplementary Figure S21**: Distribution performances for explored modes evaluated by type of encoder

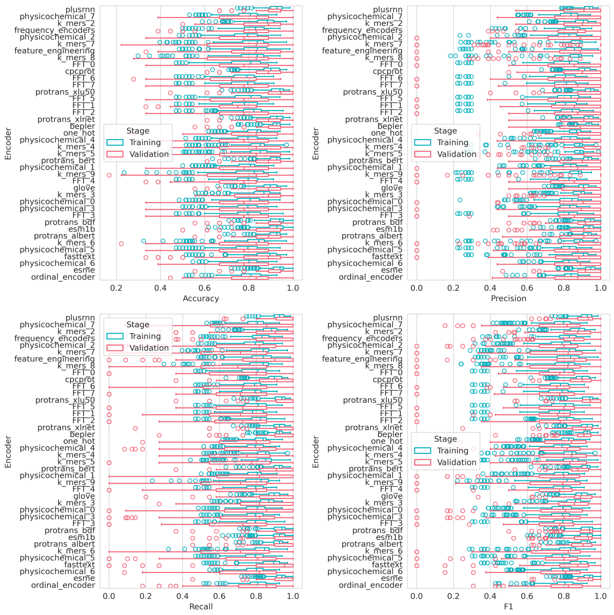


**Supplementary Figure S22**: Distribution performances for explored modes evaluated by encoder

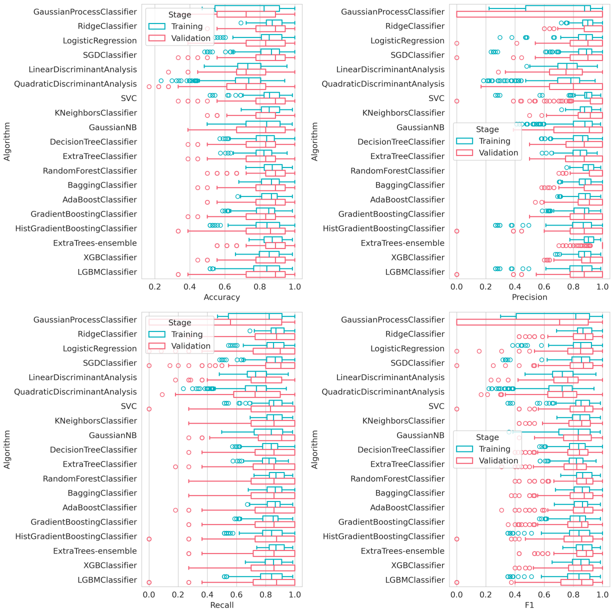


**Supplementary Figure S23**: Distribution performances for explored modes evaluated by algorithm

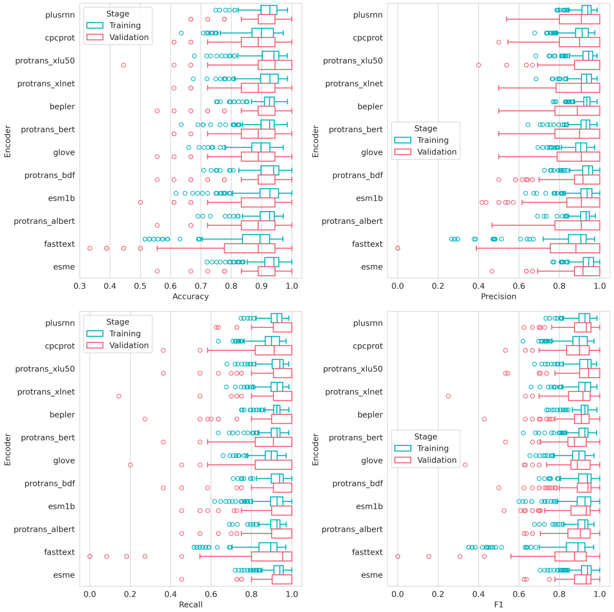


**Supplementary Figure S24**: Distribution performances for explored modes evaluated by type of pre-trained model

#### S3.7 Classification models for PU/PUR

Figure **S25** summarizes the performances obtained during explorations strategies by type of encoder approaches. Figure **S26** summarizes the performances by encoder strategy. Figure **S27** summarizes the performances by supervised learning algorithms, and Figure **S28** summarizes the performances by type of pre-trained protein language models (embedding).

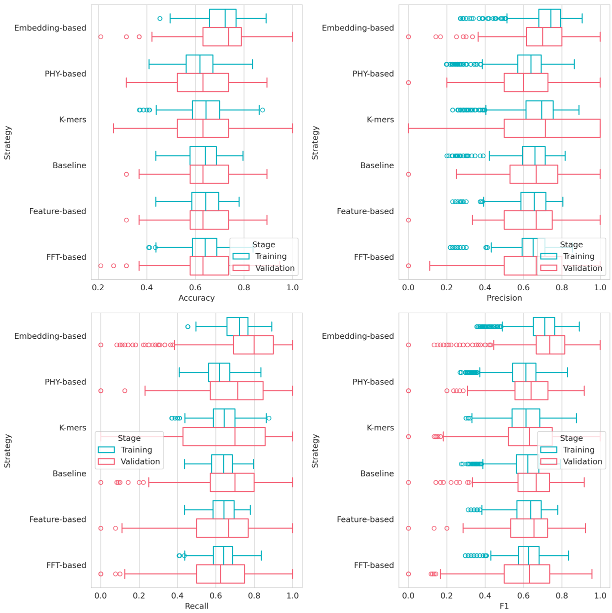


**Supplementary Figure S25**: Distribution performances for explored modes evaluated by type of encoder

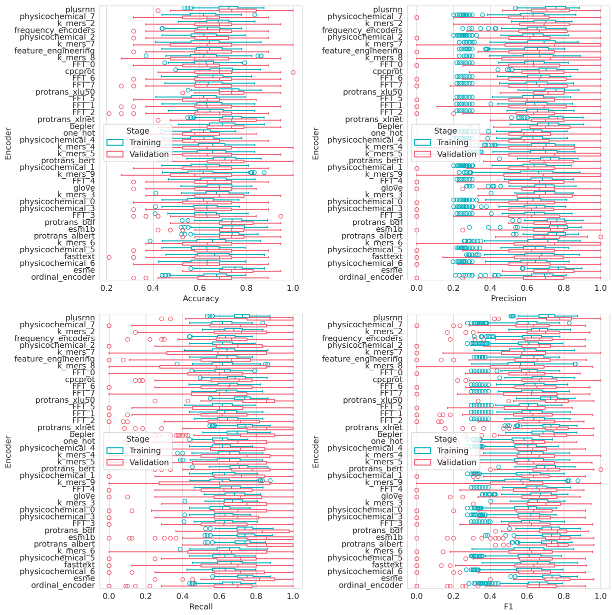


**Supplementary Figure S26**: Distribution performances for explored modes evaluated by encoder

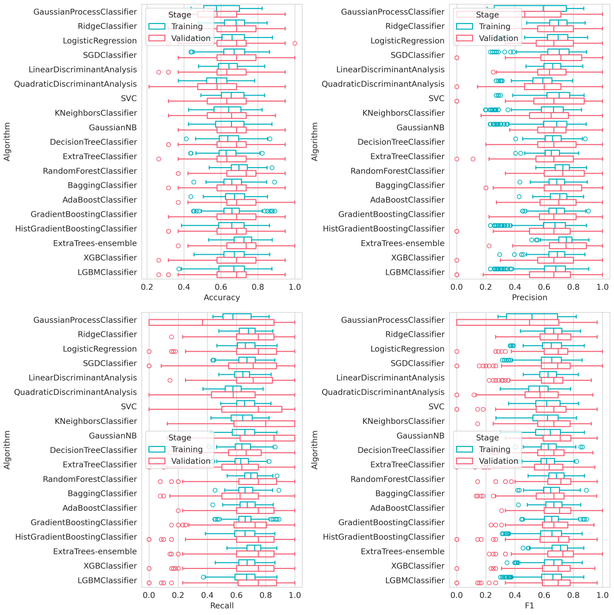


**Supplementary Figure S27**: Distribution performances for explored modes evaluated by algorithm

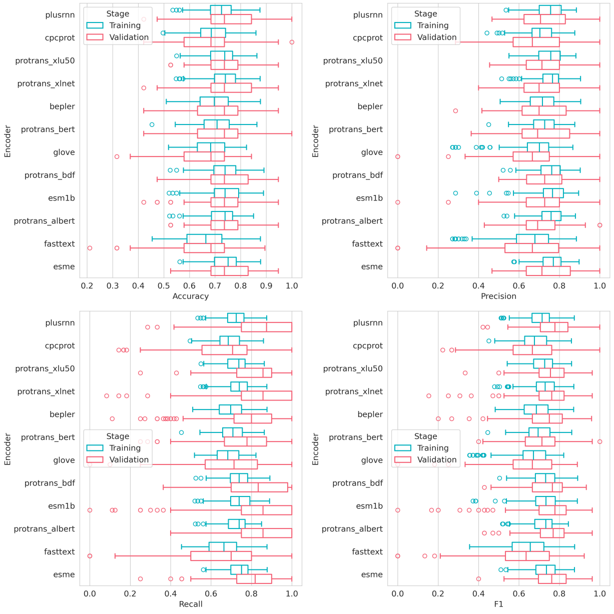


**Supplementary Figure S28**: Distribution performances for explored modes evaluated by type of pre-trained model

#### S3.8 General analysis of explored strategies

Figure **S29** summarize the F1-score distribution obtained during the exploration stage by the type of plastic evaluated in this work. Figure **S30** shows the F1-score distribution by numerical representation strategies evaluated in this work for all plastic-substrate targets explored. Figure **S31** illustrates de F1-score by embedding-based representation during the exploration stage, and Figure **S32** summarizes the F1-score by algorithms considering only embedding-based strategy.

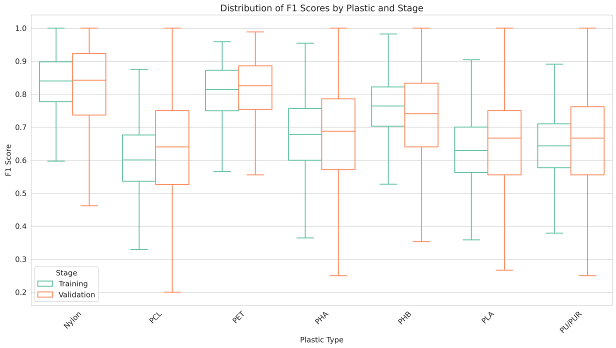


**Supplementary Figure S29**: F1-score distribution by type of plastic analyzing during training and validation process in the exploration step of the development of classification models.

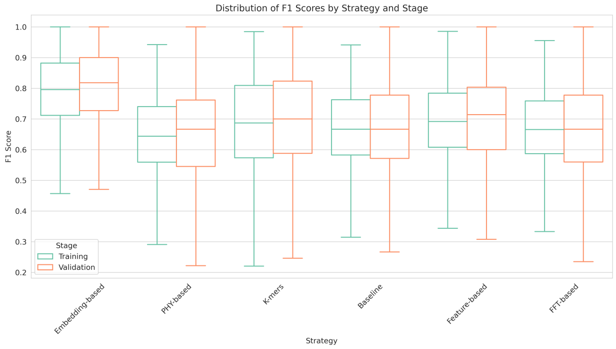


**Supplementary Figure S30**: F1-score distribution by type numerical representation strategy analyzing during training and validation process in the exploration step of the development of classification models.

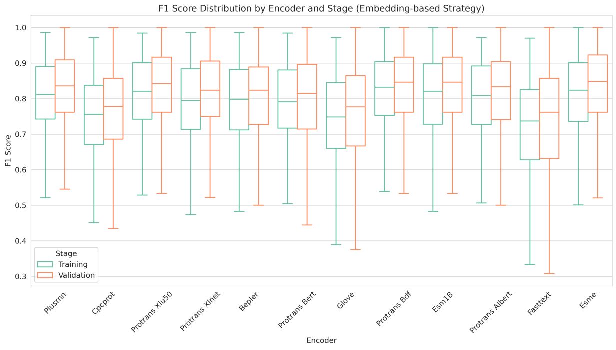


**Supplementary Figure S31**: F1-score distribution by type pre-trained model strategy analyzing during training and validation process in the exploration step of the development of classification models.

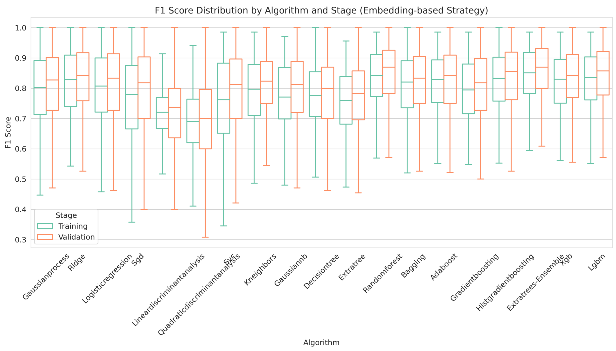


**Supplementary Figure S32**: F1-score distribution by supervised learning algorithm filter by only embedding-based numerical representation strategy analyzing during training and validation process in the exploration step of the development of classification models.

#### S4 Evaluation of generated sequences through generative learning strategies

This work generated more than 100,000 enzyme sequences by applying generative learning strategies. Using the classification models, more than 2,900 enzymes were classified as PET-degrading. The generated sequences were compared with the experimentally validated sequences using the frequency amino acid composition (see Figure **S33**) and nine physicochemical properties (see Figure **S34**).

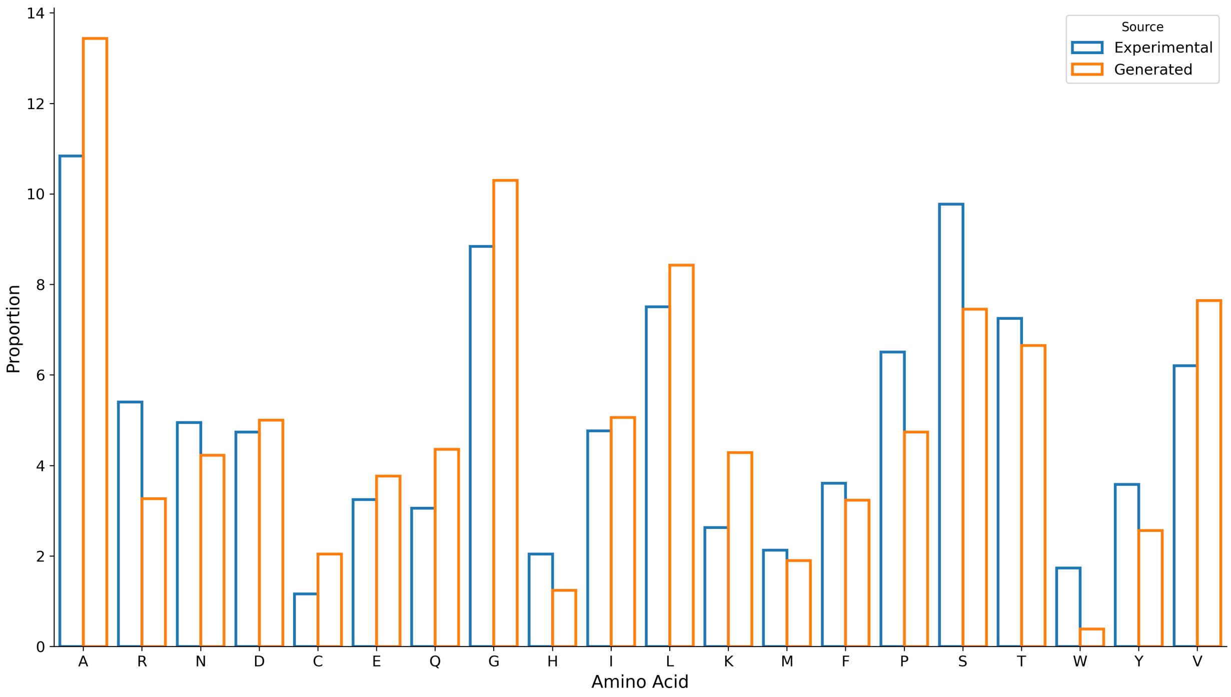


**Supplementary Figure S33**: amino acid frequency comparison between generated and experimentally validated PET-degrading enzymes.

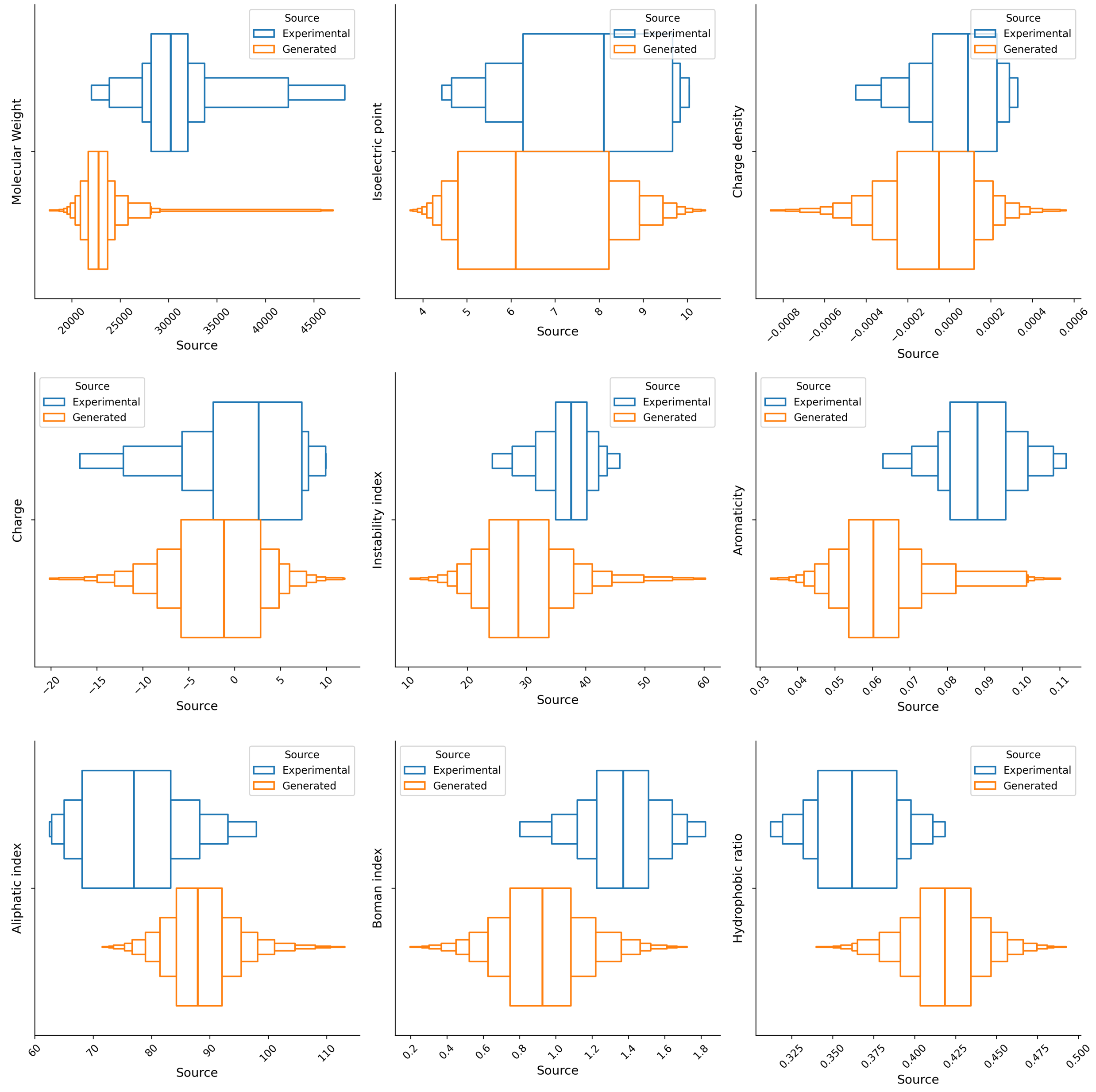


**Supplementary Figure S34**: Physicochemical distribution comparison between generated PET-degrading enzymes and experimentally validated enzymes.
